# Single-Molecule Direct RNA Sequencing Reveals the Shaping of Epitranscriptome Across Multiple Species

**DOI:** 10.1101/2023.11.16.567334

**Authors:** Ying-Yuan Xie, Zhen-Dong Zhong, Hong-Xuan Chen, Yuan-Tao Qiu, Ze-Hui Ren, Ye-Lin Lan, Fu Wu, Jin-Wen Kong, Ru-Jia Luo, Delong Zhang, Biao-Di Liu, Yang Shu, Feng Yin, Jian Wu, Zigang Li, Zhang Zhang, Guan-Zheng Luo

## Abstract

N6-methyladenosine (m6A) is an essential RNA modification that regulates gene expression and influences diverse cellular processes. Yet, fully characterizing its transcriptome-wide landscape and biogenesis mechanisms remains challenging. Traditional next-generation sequencing (NGS) methods rely on short-reads aggregation, overlooking the inherent heterogeneity of RNA transcripts. Third-generation sequencing (TGS) platforms offer direct RNA sequencing (DRS) at the resolution of individual RNA molecules, enabling simultaneous detection of RNA modifications and RNA processing events. In this study, we introduce SingleMod, a deep learning model tailored for precise m6A modification mapping on individual RNA molecules from DRS data. Applying SingleMod to human cell lines, we systematically dissect the transcriptome-wide m6A landscape at single-molecule and single-base resolution, characterizing m6A heterogeneity in RNA molecules from the same transcript and revealing that multiple m6A sites on an RNA molecule can cumulatively influence its splicing and stability. Through comparative analyses across eight diverse species, we quantitatively elucidate three distinct m6A distribution patterns that suggest divergent regulatory mechanisms. This study provides a novel framework for understanding the shaping of epitranscriptome in a single-molecule perspective.

## Introduction

Extensive researches have highlighted m6A as the predominant RNA modification in eukaryotic mRNA^1,2^, exerting substantial influence over a broad spectrum of physiological and pathological processes due to its crucial role in regulating gene expression^3–5^. Advanced mapping techniques have been devised to map m6A sites across various tissues and cell types^6^, unveiling key features of m6A deposition, such as a pronounced enrichment surrounding the stop codon and the presence of the consensus sequence DRACH in mammals^7,8,9^. Recent studies have further implicated the exon junction complex (EJC) in modulating m6A deposition through a spatial exclusion mechanism^10–13^. However, a comprehensive understanding of how m6A interacts with other RNA processing events and how these mechanisms collectively shape the observed m6A landscape remains elusive. Moreover, the extent to which these mechanisms are conserved across species is largely unknown.

Current high-throughput m6A mapping methods predominantly relies on next-generation sequencing (NGS)^7,8,14–18^, which involves analyzing short cDNA fragments generated through RNA fragmentation, reverse transcription, and PCR amplification. These steps inherently lead to the loss of crucial information about the native RNA molecules, including precise location of m6A modifications within full-length transcripts, the heterogeneity of modifications among individual transcripts, and the complex interplay between m6A and pre-RNA processing events such as alternative splicing (AS) and alternative polyadenylation (APA). Consequently, conventional NGS-based approaches are limited in their ability to fully decipher the complexity of the native epitranscriptome, the complete set of RNA molecules within a cell.

Direct RNA sequencing (DRS) technique provided by Oxford Nanopore Technology (ONT) platform has revolutionized the field by enabling the direct sequencing of native RNA molecules, offering the potential to detect RNA modifications, particularly m6A, along full-length transcripts^19,20,21^. Despite its potential, most existing computational tools designed for predicting m6A from DRS data are limited to site-level modification status and lack the resolution to determine modification on individual RNA molecules^21^. This limitation primarily arises from the lack of standardized single-molecule m6A labels, with tool development often relying on biased synthesized data or non-quantitative site-level labels from one organism, resulting in reduced accuracy and limited generalization capacity^22–26^. While recent tools like m6Anet have introduced sophisticated algorithms, such as multiple instance learning, to improve site-level m6A prediction and provide probabilities of read-level modification^21,27^, they are not designed to fully utilize the absolute quantification information provided by the latest “gold-standard” NGS-based methods^14^. Therefore, there is still a pressing need for tools that enable accurate mapping of m6A modifications at the single-molecule level from DRS data.

Here, we introduce SingleMod, a deep learning model designed for the precise detection of m6A within individual RNA molecules from nanopore DRS data. SingleMod employs a deep multiple instance regression (MIR) framework, leveraging extensive methylation-rate labels provided by the absolutely quantitative NGS-based method^14^. We demonstrate that SingleMod significantly outperforms existing tools in accuracy and generalizability across multiple datasets. Applying SingleMod to human cell lines, we explore the epitranscriptome from a single-molecule and single-base perspective, revealing that despite m6A modifications introducing substantial heterogeneity among RNA molecules, multiple m6A sites collectively shape the epitranscriptome in a position-independent, additive mode. We further demonstrate that m6A has a modest influence on AS and APA, while these processes, in turn, extensively affect m6A deposition. Furthermore, by conducting a comparative analysis of single-molecule m6A profiles across eight species with distant phylogenetic relationship, including mammals, fish, plants, protozoa, and algae, we quantitatively delineate three distinct m6A distribution patterns. Notably, we observe a vertebrate-specific pattern in line with the EJC-mediated exclusion deposition model previously described in mammals. This study not only enhances our understanding of m6A modification but also provides a novel framework for investigating the complexity of epitranscriptome in a new single-base and single-molecule perspective across diverse species.

## Result

### Development of SingleMod for single-molecule m6A detection

To develop a tool for detecting m6A at single-molecule level from DRS, we initially planned to train a conventional classification model utilizing fully-methylated (i.e., all RNA molecules at that site are modified) and -unmethylated (i.e., no RNA molecules at that site are modified) m6A sites identified by the most reliable NGS-based method^14^ to generate positive and negative samples, respectively. However, we noticed that the majority of m6A sites in human exhibit low methylation rates, while the fully-methylated m6A sites are very limited (Extended Data Fig. 1a). The 2-bp sequence flanking the motif also influences the current shift during nanopore sequencing, thus further increasing the sequence complexity needed for training data (Supplementary Fig. 1a). Therefore, it became evident that the limited availability of fully-methylated m6A sites and sequence diversity made training a conventional classification model unfeasible. To effectively utilize the abundant m6A sites with varying methylation rates, we developed a deep multiple instance regression (MIR) framework (Fig. 1a), which allows for the inclusion of m6A sites with a wide range of methylation rates during training. Employing a deep neural network, this framework predicts the likelihood of each read being modified at a specific site and optimizes the network’s parameters by minimizing the mean squared error (MSE) between the methylation rates calculated from the aggregated predicted statuses and benchmarks.

**Fig. 1.**
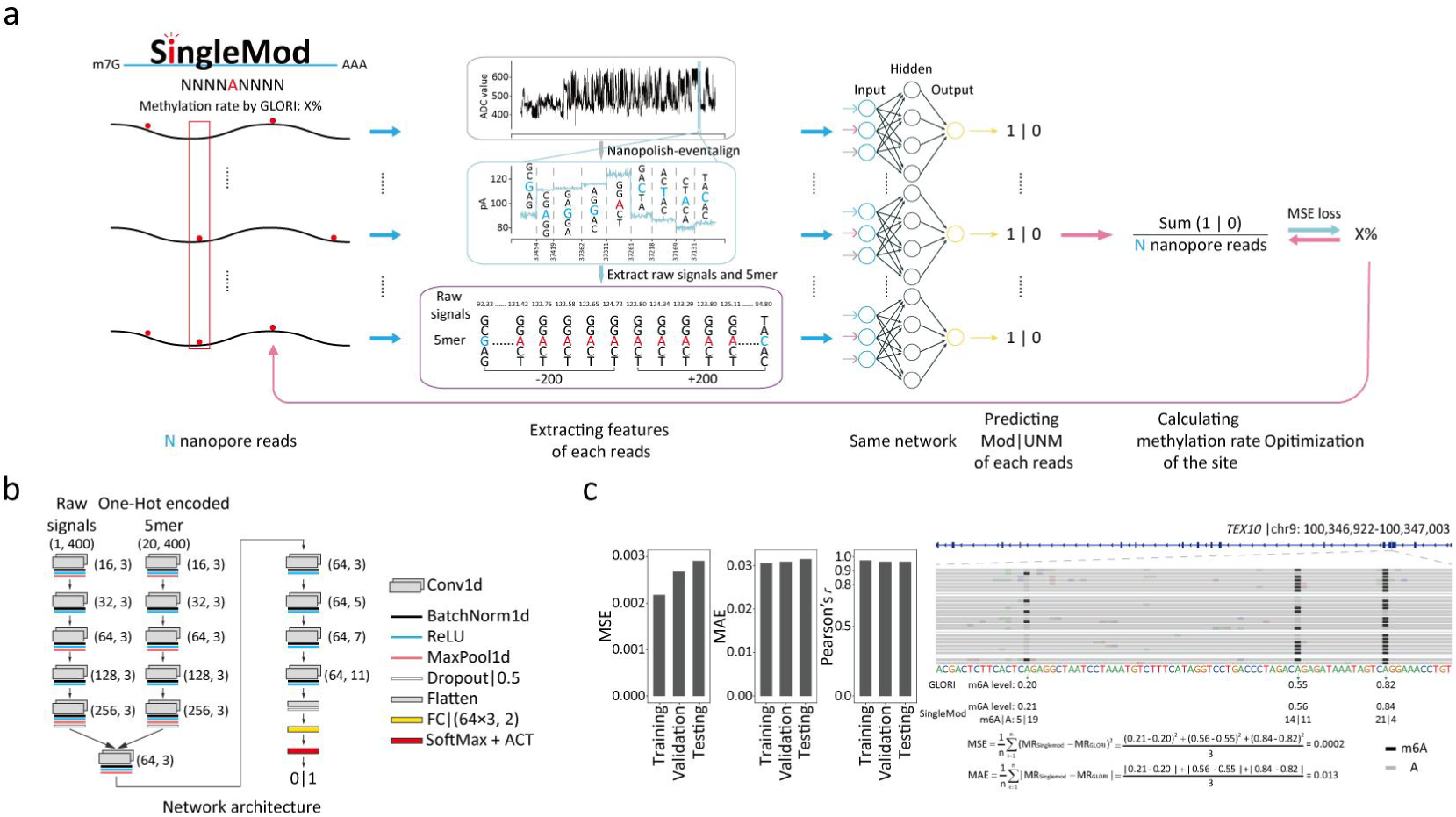
The development and training of SingleMod. **a** Methodological overview of SingleMod. **b** Architecture of the deep learning networks within the SingleMod model. Input: (channels, length), Conv1d: (output_channels, kernel_size), FC (Fully-connected): (in_features, out_features). **c** Metrics used to evaluate SingleMod, and its actual performance on the training, validation and testing sets. An IGV snapshot illustrates the single-molecule level m6A predictions and the calculation process of MSE and MAE.

To ascertain the optimal input strategy for the model, we compared its performance using two distinct types of input: one utilizing informative features extracted from raw signals, and the other directly employing raw signals. Before this comparison, we conducted incremental feature selection to search for an optimal combination of features (Supplementary Fig. 1b). Additionally, we determined the optimal length of raw signals (Supplementary Fig. 1c). We found that directly using raw signals as model input significantly outperformed using informative features (Extended Data Fig. 1b). When comparing various model architectures, we found that our custom network and the inception network (GoogLeNet-like) exhibited the best performance (Extended Data Fig. 1b). However, the inception network demonstrated slightly inferior performance with much slower convergence when implemented in MIR framework (Extended Data Fig. 1c). We therefore chose the custom network as the model architecture. Specifically, 200 flanking raw signals and corresponding One-Hot encoded 5-mer sequences are individually fed into 5-layers one-dimensional convolutions (Fig. 1a-b). Their outputs are then merged and further processed through another 5-layers one-dimensional convolution, yielding the probability of m6A modification (Fig. 1a-b). Notably, despite a significant proportion of sites exhibiting low methylation rates (Supplementary Fig. 1d), including all sites for model training did not compromise its performance for high-methylated sites; instead, it substantially improved the accuracy for low-methylated sites (Extended Data Fig. 1d-e). As a result, we included all sites for model training.

We initially trained the SingleMod model for 13 predominant motifs separately, using DRS data and methylation-rate labels derived from two human cell lines (HEK293T and HeLa) (Supplementary Fig. 1d). During the training process, SingleMod achieved satisfactory evaluation metrics in both the training and validation sets (MSE: 0.0022/0.0027, mean absolute error (MAE): 0.0306/0.0309, Pearson’s correlation coefficient (r): 0.9738/0.9657) (Fig. 1c and Supplementary Fig. 2a). Importantly, SingleMod exhibited consistent performance on the testing set, with the methylation rates of sites calculated from the aggregation of single-molecule m6A predictions closely matching the benchmarks (MSE: 0.0029, MAE: 0.0315, Pearson’s r: 0.9652) (Fig. 1c and Supplementary Fig. 2a). SingleMod not only demonstrated outstanding overall performance, but also attained comparable performance across most motifs (Supplementary Fig. 2b-d). These results demonstrated that SingleMod was successfully trained and validated.

### SingleMod accurately and robustly predicts m6A modifications at single-molecule level

To evaluate the performance of SingleMod and conduct a comparative analysis with existing tools, we implemented these tools on DRS data from two species with a distant phylogenetic relationship, namely mouse (*M. musculus*, embryonic stem cells (mESC)) and Arabidopsis (*A. thaliana*, seedlings). The results were subjected to rigorous benchmarking using two orthogonal NGS-based quantitative methods GLORI^14^ and eTAM-seq^18^. We first computed the methylation rates of sites based on the aggregation of single-molecule m6A predictions and compared them to benchmarks. To ensure a fair comparison, we focused on sites within 10 common motifs that had a methylation rate exceeding 0.1, as determined by either NGS-based methods or DRS tools. We then calculated evaluation metrics, including MSE, MAE and Pearson’s r, using these selected sites. Our results underscored the superior performance of SingleMod in both species, regardless of the choice of NGS-based methods for evaluation (Fig. 2a-b and Supplementary Fig. 4a, 5a). These superior performances were consistent across both overall assessments and individual motifs (Extended Data Fig. 2a-b and Supplementary Fig. 3, 4b, 5c). Additionally, SingleMod accurately reflected changes in methylation rates following m6A writer knockout (KO) (Fig. 2c and Supplementary Fig. 5b), which aligns with the nearly absent m6A signal in KO mESC and 10% residual in Arabidopsis mutant, as determined by LC-MS/MS^21,28^.

**Fig. 2.**
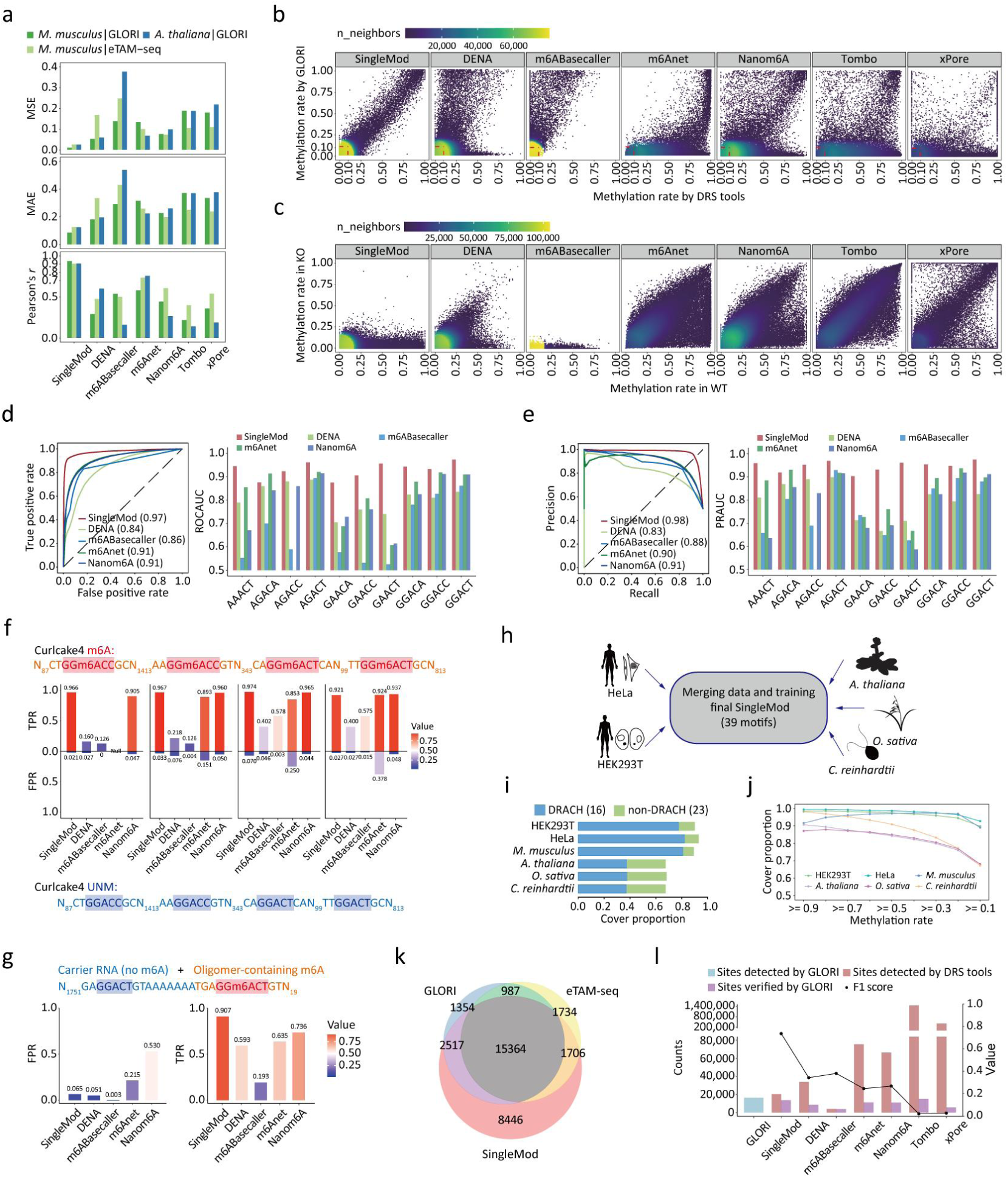
SingleMod’s accurate and robust performance across diverse datasets. **a** Comparing the overall performance on *M. musculus* and *A. thaliana* between SingleMod and other tools, using three evaluation metrics as evaluated by two NGS-Based methods. Data used in **b-e**, **k-l** were from *M. musculus*. **b** Comparing the methylation rates predicted by DRS tools to GLORI benchmarks. **c** Comparing the methylation rates between WT and *mettl3* KO mESC predicted by DRS tools. **d-e** ROC (**d**) and PR (**e**) curves for single-molecule m6A predictions within a representative motif GGACT. The AUC of ROC (**d**) and PR (**e**) curves of 10 common motifs were also provided. The AUC values of AGACC motif of m6Anet were unavailable due to a lack of sufficient data for plotting. Fully-methylated and -unmethylated sites determined by eTAM-seq were used in **d-e**. **f-g** Comparing the TPR and FPR of single-molecule level m6A predictions between SingleMod and other tools, using DRS data from the IVT-derived (Curlcakes, **f**) or synthetic (Oligomer, **g**) RNA molecules. **h** Merging data from multiple species and training the final version of SingleMod, which supports m6A prediction within 16 DRACH and 23 non-DRACH motifs. **i** Proportion of m6A sites (methylation rate >= 0.1 as determined by GLORI) falling within SingleMod’s motifs scope. **j** Proportion of m6A sites with varying methylation rates as determined by GLORI falling within SingleMod’s motifs scope. **k** Overlap of identified m6A sites within the candidate list by SingleMod, GLORI, and eTAM-Seq. **l** The number of identified m6A sites within the candidate list by DRS tools and GLORI, and their overlap. F1 scores (a single metric combines Precision and Recall) of site-level m6A predictions were also provided. Tombo and xPore were not included in **d-g**, as they cannot produce single-molecule level m6A predictions.

To directly evaluate the accuracy of single-molecule m6A predictions, we selected fully-methylated (methylation rate > 0.99) and -unmethylated (methylation rate < 0.01) sites determined by GLORI or eTAM-seq, and assigned the labels of corresponding molecules from these sites as methylated (1) and unmethylated (0), respectively. Comparing to other tools, SingleMod achieved the best area under curve (AUC) of receiver operating characteristic (ROC) and precision recall (PR) curve in 9 out of 10 motifs and maintained robust performance across most motifs (Fig. 2d-e and Supplementary Fig. 5d). We further applied SingleMod to DRS data from three types of synthetic RNA molecules with m6A (or A) introduced at specific locations through either in vitro transcription (IVT) or chemical synthesis (Curlcakes^20^, Oligomer^23^ and SL-Oligomers^29^, see methods). SingleMod demonstrated superior sensitivity in detecting multiple m6A modifications on individual RNA molecules, alongside particularly well-balanced true positive rates (TPR) and false positive rates (FPR) (Fig. 2f-g and Extended Data Fig. 2c-d). Across the majority of sites, it achieved a sensitivity (reflected by TPR) of nearing 96% and a specificity (reflected by FPR) of exceeding 97% (Fig. 2f-g and Extended Data Fig. 2c-d). Notably, the latest tool m6ABasecaller effectively controlled FPR, while compromising its TPR (Fig. 2f-g and Extended Data Fig. 2c). Nanom6A, originally trained on IVT-derived RNA, naturally performed well on this dataset but exhibited significantly diminished accuracy on other synthetic RNAs (Fig. 2f-g, Extended Data Fig. 2c). In conclusion, SingleMod demonstrated exceptional accuracy and robustness at single-molecule level, surpassing the performance of any other existing tools.

### Enhancing generalization by integrating data from species with distant phylogenetic relationship

We noticed that m6A is predominantly enriched within several DRACH motifs in mammals, while spreading in a broader spectrum of motifs in plants and other eukaryotic organisms (Supplementary Fig. 6a). Furthermore, certain high-methylated sites were underestimated by SingleMod, which may occur within species-specific sequence contexts that have not been sufficiently trained (Supplementary Fig. 6b-c). Therefore, we hypothesized that training using data from a single species might introduce biases due to limited motifs and sequence diversity, potentially limiting the model’s generalization ability. To address this, we combined datasets from human (HEK293T, HeLa), plants (*A. thaliana, O. sativa*) and green algae (*C. reinhardtii*) to re-train the models (Fig. 2h). By integrating data from species with distant phylogenetic relationship, we obtained sufficient training data for up to 16 DRACH motifs and 23 non-DRACH motifs, collectively covering approximately 90% of known m6A sites in mammals and about 70% in plants and green algae (Fig. 2i). Importantly, this coverage for m6A sites with higher methylation rates is notably increased (Fig. 2j). This integration also enabled us to utilize m6A sites from a more diverse sequence context to train our models (Supplementary Fig. 6d).

To evaluate the improved generalization, we applied the final version of SingleMod to DRS data from mouse, assessing whether it could recall the majority of m6A sites while maintaining high precision. Comparing to other tools, the results of SingleMod exhibited substantial overlap with those derived from two NGS-based methods. Specifically, SingleMod achieved a considerably superior F1 score, with a recall of 0.82 and a precision of 0.67 (Fig. 2k-l and Supplementary Fig. 4c). Therefore, this strategy substantially enhanced the generalization of the model when applied to other species (Supplementary Table 4). To make SingleMod compatible with the latest sequencing kit (RNA004), we also used the new DRS data to train our model (Supplementary Fig. 7a-h and Supplementary Table 4). Notably, SingleMod consistently outperformed ONT’s official tool, Dorado, across all available motifs, while Dorado introduced extensive false positives beyond the canonical DRACH motifs (Supplementary Fig. 8).

### Single-molecule m6A landscape in human cell lines

Leveraging the high accuracy of SingleMod in detecting m6A at single-molecule resolution, we depicted the m6A landscapes in three human cell lines (HEK293T, HeLa, and K562) and analyzed the genomic features from a single-molecule perspective (Fig. 3a-b). Strikingly, we found that approximately 50% of the mRNA molecules lack m6A modifications in all cell lines (Fig. 3c), contrasting with previous findings obtained from site-level m6A mapping methods^30^. However, further analysis unveiled that these unmodified molecules are primarily transcribed from a small subset of highly expressed genes, such as those coding for ribosomal proteins (Extended Data Fig. 3a-b). For the majority of genes or isoforms, a significant proportion of their corresponding RNA molecules have at least one m6A modification (Fig. 3d and Supplementary Fig. 9d, j). For instance, over 54% of genes in HEK293T demonstrate that more than 90% of their RNA molecules possess m6A modifications (Fig. 3d).

**Fig. 3.**
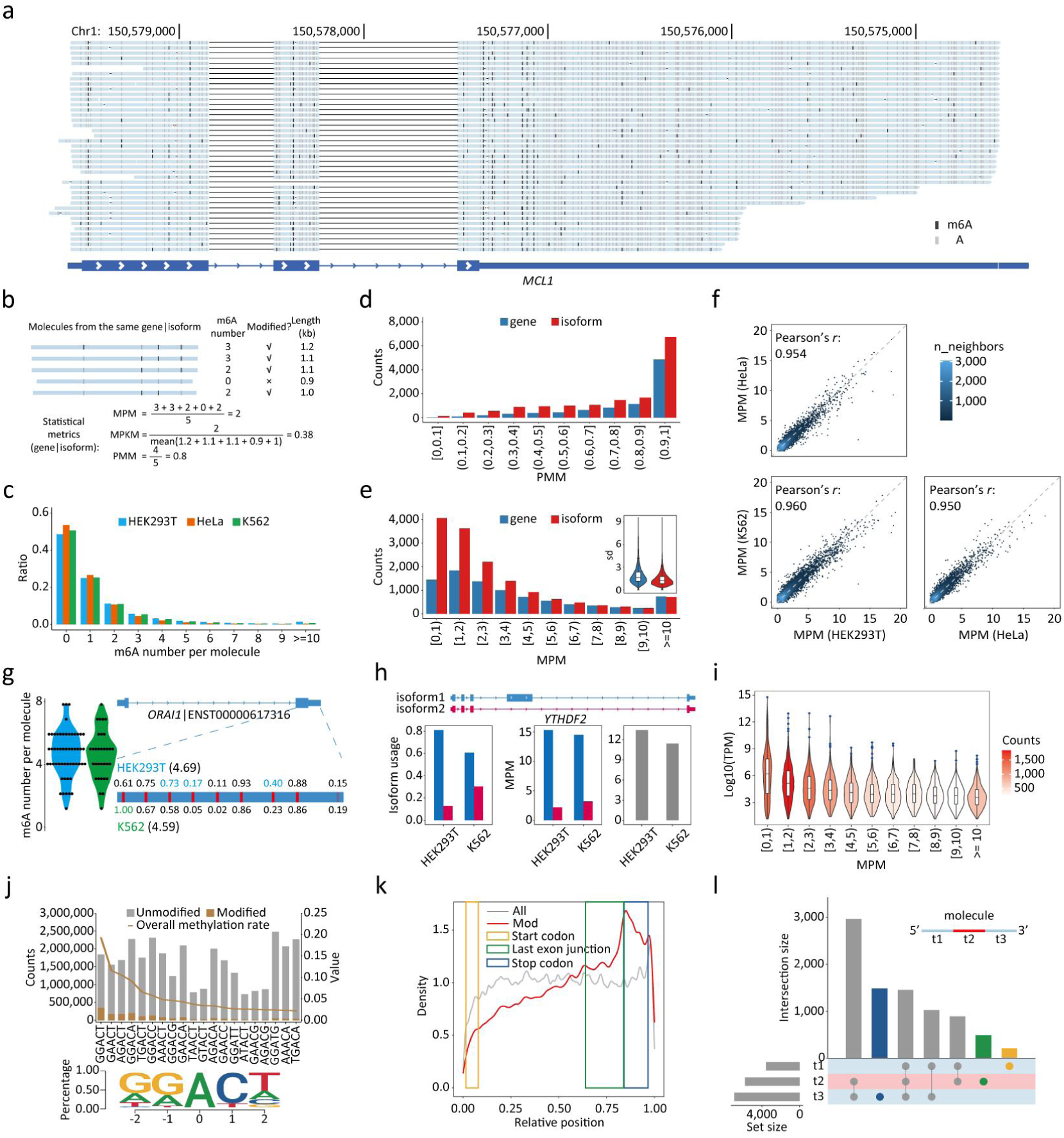
Single-molecule m6A profiling in human cell lines by SingleMod. **a** A representative IGV snapshot illustrates the single-molecule m6A predictions by SingleMod for gene *MCL1.* The predicted status of each potential site on each individual molecule was indicated. **b** Schematic illustration depicts the calculation of single-molecule level m6A quantification metrics for genes or isoforms. The MPM represents the average number of m6A modifications per RNA molecule, the MPKM represents the average number of m6A modifications per RNA molecule per 1,000 bp length, and the PMM represents proportion of modified molecules (with at least one m6A modification). **c** Ratio of molecules with varying number of m6A in three human cell lines. **d** Counts of genes or isoforms with varying PMM. Data used in **d-e**, **i-l** were from HEK293T. **e** Counts of genes or isoforms with varying MPM, and the distribution of standard deviation in m6A number among molecules within each gene or isoform. **f** Comparison of the MPM for identical isoforms across different cell lines. **g** Comparison of the distribution of m6A number in all molecules of a representative isoform (left), the MPM of this isoform, and the methylation rates of each site (right) between HEK293T and K562. **h** Comparison of the isoform usages within a representative gene (left), the MPM of these isoforms (middle), and the MPM of this gene (right) between HEK293T and K562. **i** Distribution of expression levels of genes with varying MPM. “TPM” indicates Transcripts Per Million. **j** Counts (top, bar) of unmodified and modified A bases from all molecules for top 20 motifs and corresponding overall methylation rates (top, line). The composition of m6A flanking bases plotted by Seqlogo was shown below. **k** The distribution of relative position of background (All) and modified (Mod) A bases within their respective molecules. The relative regions of the start codon, last exon junction and stop codon were also marked. The relative position is defined as the ratio of the distance from the m6A site to the 5’ end of the molecule to the total length of the molecule. **l** Upset plot shows the counts of gene with different m6A distribution patterns. RNA molecules were divided into three equal intervals, focusing on the difference in distribution of m6A relative position among genes. For a specific gene, if over 25% of m6A modifications fall within one interval, it is deemed to have significant m6A modifications in that interval.

For a more in-depth quantitative exploration, we introduced two metrics to characterize m6A levels for each gene or isoform. These metrics calculate the average number of m6A modifications per RNA molecule (MPM), or per RNA molecule per 1,000 bp length (MPKM), which reflect the absolute m6A quantity and density, respectively (Fig. 3b). In HEK293T, though approximately 70% of genes or isoforms exhibit an MPKM below two, 63% and 49% of them display an MPM of two or more (Fig. 3e and Extended Data Fig. 3c). These values are quite similar in K562 and slightly different in HeLa, where the proportion of high-methylated genes or isoforms is relatively lower (Supplementary Fig. 9e, k). Notably, the heterogeneity in m6A numbers at the gene level is greater than that at the isoform level, as indicated by the larger standard deviation among distinct RNA molecules (Fig. 3e, Extended Data Fig. 3d and Supplementary Fig. 9e, k). This highlights the importance of single-molecule m6A mapping, as gene-level quantification may not accurately represent m6A levels of its isoforms (Extended Data Fig. 3e).

While previous studies using site-level m6A quantification reported considerable variation in m6A levels across different cell lines or tissues, our single-molecule m6A quantification revealed that the MPM for the vast majority of isoforms is remarkably consistent among the three cell lines (Fig. 3f), primarily due to infrequent occurrence of multiple m6A sites within the same isoform exhibiting concordant changes in methylation levels (Fig. 3g and Extended Data Fig. 3f). Nonetheless, differential isoform usage may contribute to variations in MPM at the gene level (Fig. 3h), and higher MPM consistently coincide with an overall decrease in expression levels (Fig. 3i and Supplementary Fig. 9a, f, l). Furthermore, the motif preference for m6A modifications exhibits high similarity among three cell lines, with GGACT, GAACT, AGACT, GGACA, and TGACT being the most frequently modified motifs (Fig. 3j and Supplementary Fig. 9g, m). However, the overall methylation rates for most of motifs in HeLa are slightly lower than those in HEK293T and K562 (Fig. 3j and Supplementary Fig. 9g, m). Following *METTL3* knockout, a general reduction in the overall methylation rate was observed for all motifs (Extended Data Fig. 3g).

Leveraging the advantages of long-read sequencing, we endeavor to redefine the distribution of m6A across full-length RNA molecules. Generally, m6A is enriched toward the 3’ end, specifically within the region of relative positions 0.6 to 1 (Fig. 3k and Supplementary Fig. 9h, n). By annotating each molecule with key mRNA landmarks, including the start codon, last junction and stop codon, we observed that the degree of m6A enrichment increases and subsequently decreases after the last junction, the last exon still contains the predominant m6A modifications (Fig. 3k and Supplementary Fig. 9b, h, n). Consistently, the proportion of modified DRACH motifs increases significantly toward the 3’ end, reaching its peak around 200 bp from the last junction or 100 bp from the stop codon and remaining stable over an extended distance (Extended Data Fig. 3h-i and Supplementary Fig. 9c, i, o). Despite the general enrichment at the 3’ end, there are distinct, gene-specific m6A distribution patterns, with a substantial portion of genes exhibiting significant m6A levels at their 5’ end and the middle regions (Fig. 3l).

### Dissecting m6A-mediated RNA heterogeneity and regulation at the single-molecule level

We next investigated the impact of m6A modifications on RNA heterogeneity and regulation at the single-molecule level. Previous studies have shown that m6A sites tend to cluster in “hot regions”^14,17^. Supporting this notion, we observed a significantly shorter distance than expected between two adjacent m6A modifications on individual molecule (Extended Data Fig. 4a). Nevertheless, within a cluster region, it remains unclear whether m6A modifications are enriched within certain RNA molecules or if it’s randomly distributed among different molecules. Our initial assessment of adjacent m6A sites demonstrated a probabilistic distribution rather than a concurrent pattern at the molecule level (Fig. 4a and Supplementary Fig. 10f, i). This suggests that the presence of m6A at a given site does not depend on the modification status of an adjacent site within a specific RNA molecule. Consistently, RNA molecules with m6A modifications numbered according to the probabilistic distribution are the most abundant, whereas those with no or complete modifications are rare (Fig. 4b and Supplementary Fig. 10g, j). Consequently, only a small proportion of RNA molecules have multiple m6A modifications located within a cluster (Extended Data Fig. 4b). Furthermore, at the gene level, the number of m6A modifications across different molecules appears normally distributed, and m6A modifications are not enriched at specific locations (Supplementary Fig. 10a-b). These findings collectively indicate that within a gene, m6A modifications are evenly distributed across multiple molecules and multiple sites, resulting in a diverse population of RNA molecules with varying m6A signatures. We quantified this molecular heterogeneity by measuring the probability of randomly selecting two identical molecules within a gene or isoform, further demonstrating that m6A modifications substantially increase the level of heterogeneity (Fig. 4c and Supplementary Fig. 10c, h, k).

**Fig. 4.**
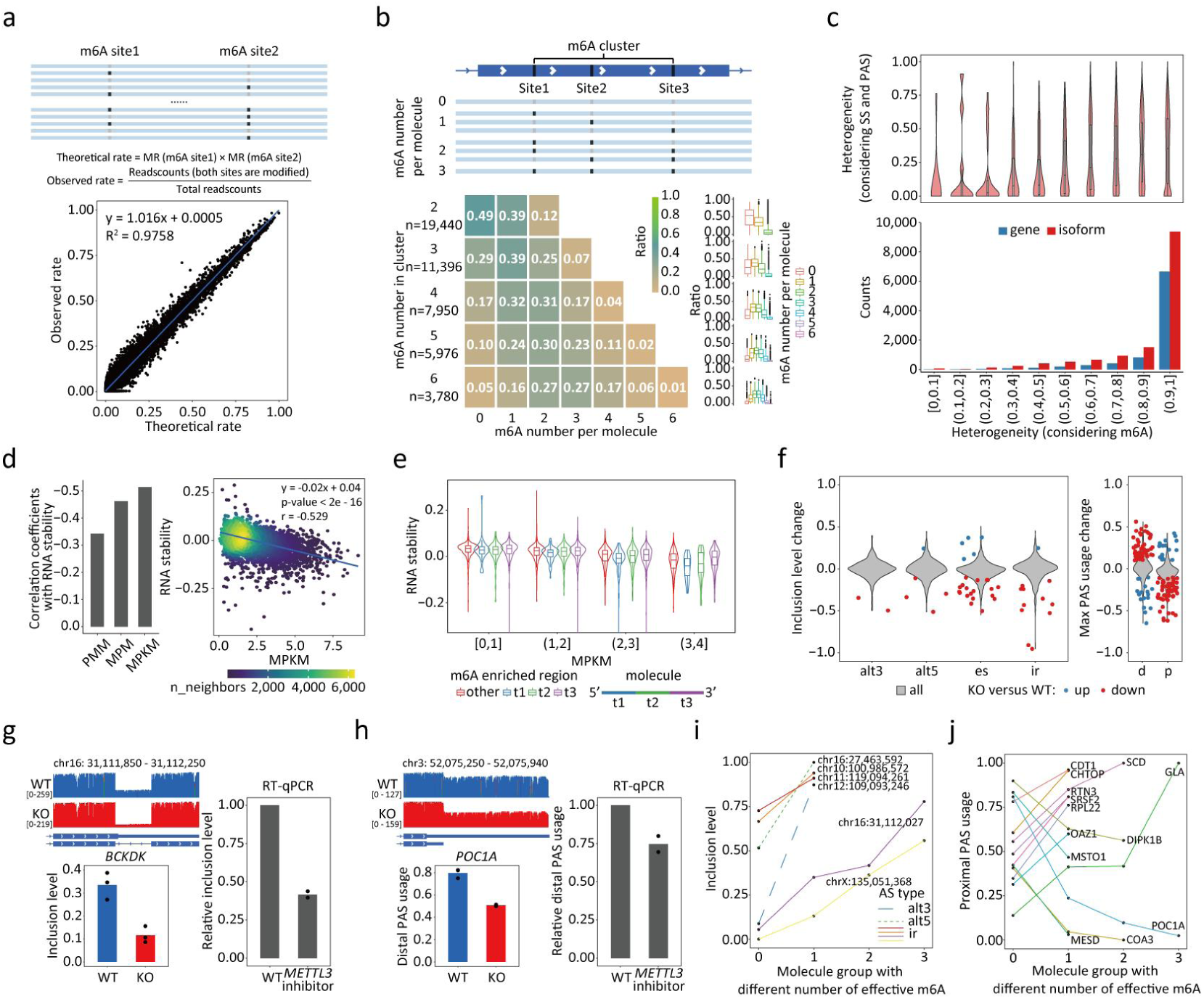
m6A-mediated RNA heterogeneity and regulation profiled at single-molecule level in human. **a** Top: schematic illustration depicts the calculation of theoretical rate and observed rate of molecules where two adjacent A sites within detected motifs are both modified. Bottom: comparison between these two values. **b** Schematic illustration (top) depicts molecules displaying possible m6A patterns within an m6A cluster. Heatmap (bottom, left) and boxplot (bottom, right) show the ratio of molecules with varying m6A number within m6A cluster. **c** Heterogeneity among RNA molecules within a gene or isoform was represented by 1 - the probability of randomly selecting two identical molecules. Top: distribution of molecular heterogeneity of corresponding genes only considering difference on SS and PAS selections. Bottom: genes or isoforms counts with varying molecular heterogeneity introduced by m6A. **d** Left: Pearson’s r between RNA stability and three m6A quantification metrics. Right: correlation between RNA stability and MPKM. **e** Distribution of RNA stability of genes with varying MPKM and m6A enriched regions. **f** Distribution of change in inclusion level of AS events (left) and max PAS usage (right) following *METTL3* KO. “alt3”, “alt5”, “es”, “ir”, “d” and “p” indicate alternative 3’ ss, alternative 5’ ss, exon skipping, intron retention, distal PAS and proximal PAS, respectively. **g-h** Two examples of differential ir (**g**) and apa (**h**) event and their corresponding validation results in cells treated with the *METTL3* inhibitor via RT-qPCR. In RT-qPCR experiment, ΔΔCt method was used to quantify the relative inclusion level (or distal PAS usage) between WT and *METTL3* inhibitor-treated cells, and this value of WT was set to 1. Two replicates were used in RT-qPCR experiment. **i-j** Inclusion level (**i**) and proximal PAS usage (**j**) of molecule groups with varying numbers of effective m6A.

The observed molecular heterogeneity prompted us to investigate the mechanisms by which m6A modifications exert their regulatory effects. The most widely recognized function of m6A is promoting RNA degradation^31,32^. Supporting this, we observed a negative correlation between the change in m6A levels and gene expression following *METTL3* knockout (Extended Data Fig. 4c). Furthermore, we directly observed a clear negative correlation between m6A levels and RNA stability, with the most notable correlation associated with the MPKM (Fig. 4d and Supplementary Fig. 10d). Importantly, the correlation between MPKM and RNA stability persists regardless of the specific positions of m6A modifications (Fig. 4e). Consistently, gene categories expected to have lower RNA stability levels, such as lncRNA, exhibit higher MPKM (Supplementary Fig. 10e). These results imply that the function of m6A in promoting RNA degradation is position-independent and operates in an additive manner.

As DRS simultaneously captures m6A and RNA processing signatures, we compared the splicing site (SS) and polyadenylation site (PAS) selections between WT and *METTL3* KO cells, aiming to further unravel the role of m6A in pre-RNA processing. Contrary to previous understanding^33,34^, the vast majority of AS events remain unchanged in KO cells, with only a small fraction showing differences, primarily a decrease in inclusion levels (Fig. 4f and Extended Data Fig. 4d). However, m6A appears to exert a stronger impact on APA, with 137 genes displaying some degree of differential PAS usage, among which 104 genes exhibit reduced usage of proximal PAS (Fig. 4f and Extended Data Fig. 4d). Although most of these differential AS and APA events show relatively modest changes, there are still some notable cases, which were further confirmed in cells treated with the *METTL3* inhibitor (Fig. 4g-h, Extended Data Fig. 4e-g and Supplementary Table 7). To identify effective m6A sites and provide molecule-level evidence for their impact on SS and PAS selection, we statistically analyzed four types of molecule species within the WT sample: those with or without m6A modification at a specific location and with or without selection of a specific SS or PAS. This analysis identified effective m6A sites for 19 differential AS and APA events, revealing that the presence of m6A modification at these sites influences the selection of SS or PAS in corresponding pre-RNA processing events (Fig. 4i-j, Extended Data Fig. 4h-i and Supplementary Fig. 11). Specifically, the most effective m6A sites are located close to the regions where AS or APA takes place, typically inhibiting proximal SS selection while promoting proximal PAS selection. Interestingly, certain events are regulated by multiple m6A sites, which appear to work additively, independently of their specific locations (Fig. 4i-j, Extended Data Fig. 4h-i and Supplementary Fig. 11).

### m6A deposition is extensively affected by pre-RNA processing

Having observed a modest impact of m6A on pre-RNA processing, we wonder whether the pre-RNA processing, in turn, affects m6A deposition. The high heterogeneity of RNA molecules with varying m6A modification and SS (or PAS) selection enabled us to conduct extensive multi-to-multi correlation analyses between each m6A site and each AS (or APA) event (Fig. 5a, h). Through this analysis, we identified numerous significantly correlated m6A-AS pairs; however, the majority of these AS events did not exhibit alterations in inclusion levels following *METTL3* knockout (Fig. 5b), suggesting that m6A deposition is affected by the process or outcome of pre-RNA splicing. Specifically, we found that RNA molecules with different SS selections exhibit distinct m6A levels at associated sites, with m6A being suppressed when a proximal SS is selected (Fig. 5c-d). This phenomenon is extensively consistent for the vast majority of AS events across different cell lines, although some do not reach statistical significance in correlation analyses (Fig. 5e and Extended Data Fig. 5a-c, i-k).

**Fig. 5.**
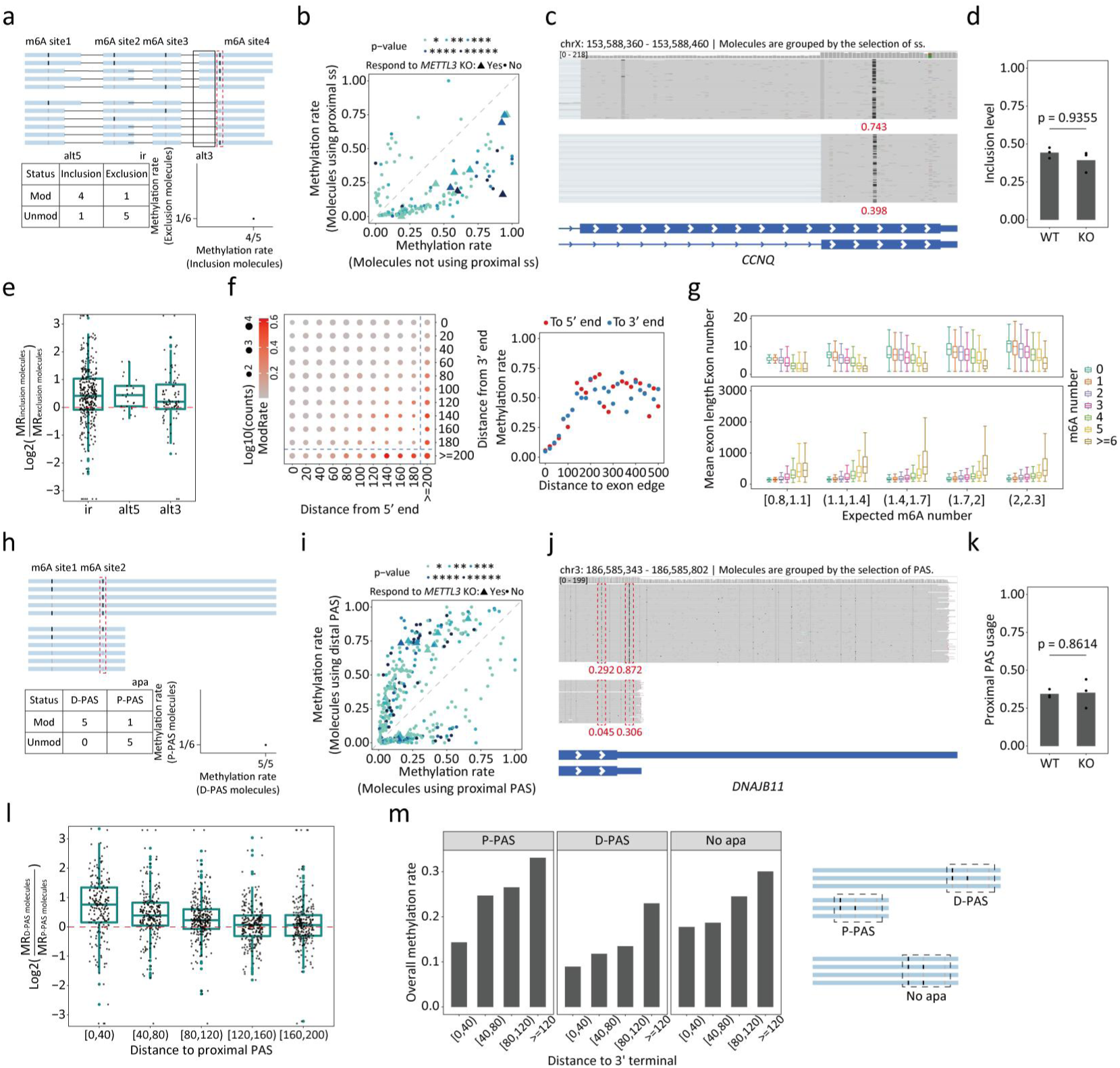
Single-molecule evidence that m6A deposition is affected by pre-RNA splicing and polyadenylation. **a** Schematic illustration depicts the molecule-level correlation analysis between m6A sites and AS events. **b** Dotplot compares the methylation rate of m6A site between two groups of molecules with different SS selections in the correlation analysis, as calculated in **a**. Each dot represents a significantly correlated m6A site-AS event pair, and its P-value and whether this correlated AS event exhibited differential splicing following *METTL3* KO were also indicated. **c-d** An IGV snapshot Illustrates a representative significantly correlated m6A site-AS event (alt3) pair. We grouped these molecules based on the 3’ SS selection and compared the methylation rates (indicated in red) of the correlated m6A site between two groups (**c**). This AS event do not show differential splicing between WT and *METTL3* KO cells (**d**). **e** Comparing the methylation rates of nearby m6A sites (<=200 bp from exon edge) between inclusion-spliced molecules and exclusion-spliced molecules across different AS events, as showed in **a**. **f** Overall methylation rates at positions with varying distance from both edges in the middle exons (left) and when controlling the distance from opposite edges >=200 (right), using GAACT motif as a representative. **g** Comparing the exon number and mean exon length of molecules with different m6A number when controlling the m6A modification potentials (the expected m6A number, considering the number and composition of motifs). **h** Schematic illustration depicts the molecule-level correlation analysis between m6A sites and APA events. D-PAS and P-PAS indicate distal and proximal PAS, respectively. **i** Dotplot compares the methylation rate of m6A site between two groups of molecules with different PAS selection in the correlation analysis, as calculated in **h**. Each dot represents a significant correlated m6A site-APA event pair, and its P-value and whether this correlated APA event exhibited differential PAS usage following *METTL3* KO were also indicated. **j-k** An IGV snapshot Illustrates a representative significantly correlated m6A site-APA event pair. We grouped these molecules based on the PAS selection and compared the methylation rates (indicated in red) of the correlated m6A site between two groups (**j**). This APA event did not show differential PAS usage between WT and *METTL3* KO cells (**k**). **l** Comparing the methylation rates of m6A sites near proximal PAS between D-PAS molecules and P-PAS molecules, as showed in **a**. **m** Overall methylation rate at positions with varying distance from the 3’ terminal of three types of molecules, using GAACT motif as a representative (left). Schematic illustration depicts the three types of molecules (right).

Recent studies have revealed that EJC, a constitutive protein complex that binds at exon-exon junctions post-splicing, spatially repels methyltransferases, thus leading to m6A suppression^10–13^. To further elucidate how EJC affects m6A, we compared the overall methylation rate at different positions categorized by their distance from the exon edges, including the 5’ end and 3’ end (Fig. 5f and Extended Data Fig. 5d, l). We found that m6A suppression effect increases as getting closer to either edge, with an influence range extending up to 150 bp from edge. This suppression effect is similar for both 5’ and 3’ end at comparable distances (Fig. 5f and Extended Data Fig. 5d, l). Notably, as most human exons are short, the majority of sites on RNA molecules fall within the influence range of EJC suppression (less than 150 bp from either edge), with cases even undergoing dual-suppression from both sides. As expected, we observed a global trend where molecules with higher number of m6A tend to undergo fewer splicing event (Fig. 5g).

In addition to AS, we identified more significantly correlated m6A-APA pairs. Similarly, m6A deposition is also affected by PAS selection, as most of these APA events did not respond to *METTL3* KO (Fig. 5i). In most cases, m6A modifications tend to be added when the RNA molecule chooses the distal PAS, although some instances associated with intronic PAS are more likely influenced by EJC (Fig. 5i-k and Extended Data Fig. 5q). Furthermore, this observation is also widespread in most APA events across different cell lines, with m6A suppression occurring within 120 bp from the proximal PAS and intensifying as it approaches the PAS (Fig. 5l and Extended Data Fig. 5e-g, m-o). We speculate that a general mechanism, independent of APA itself, may exist whereby m6A deposition is suppressed near the PAS of RNA molecules. Supporting this hypothesis, we observed that the overall methylation rates near the PAS are lower than those in other regions (≥120 bp from PAS), regardless of APA occurrence (Fig. 5m and Extended Data Fig. 5h, p).

### Comparative analysis of single-molecule m6A profiles across multiple species reveals three distinct m6A distribution patterns

The ability to map m6A at single-molecule and single-base resolution has empowered us to conduct a detailed comparative analysis of m6A profiles across various species in a new perspective. We employed SingleMod to analyze species with distant phylogenetic relationship, including mammals (*H. sapiens* and *M. musculus*), fish (*D. rerio*), plants (*A. thaliana*, *O. sativa* and *P. trichocarpa*), green algae (*C. reinhardtii*), protozoa (*T. gondii*), and nematode (*C. elegans*). Intriguingly, we found that m6A displays diverse motif preferences across different species, and this divergence clusters them into distinct categories that closely correspond to their evolutionary relationships (Fig. 6a). The composition of flanking sequences surrounding m6A is also highly consistent within each cluster (Extended Data Fig. 6a and Supplementary Fig. 12a). Additionally, different species display varying distributions of m6A numbers within single RNA molecules and MPM across isoforms and genes (Fig. 6b-c and Supplementary Fig. 12b). For example, in protozoa and zebrafish, RNA molecules exhibit a higher number of m6A modifications, which may be attributed to their longer lengths and unique motif compositions (Extended Data Fig. 6b and Supplementary Fig. 12c). The nematode was excluded from subsequent analyses due to extremely low m6A levels in all motifs, aligning with previous findings that suggest the absence of m6A in nematode mRNA^35^.

**Fig. 6.**
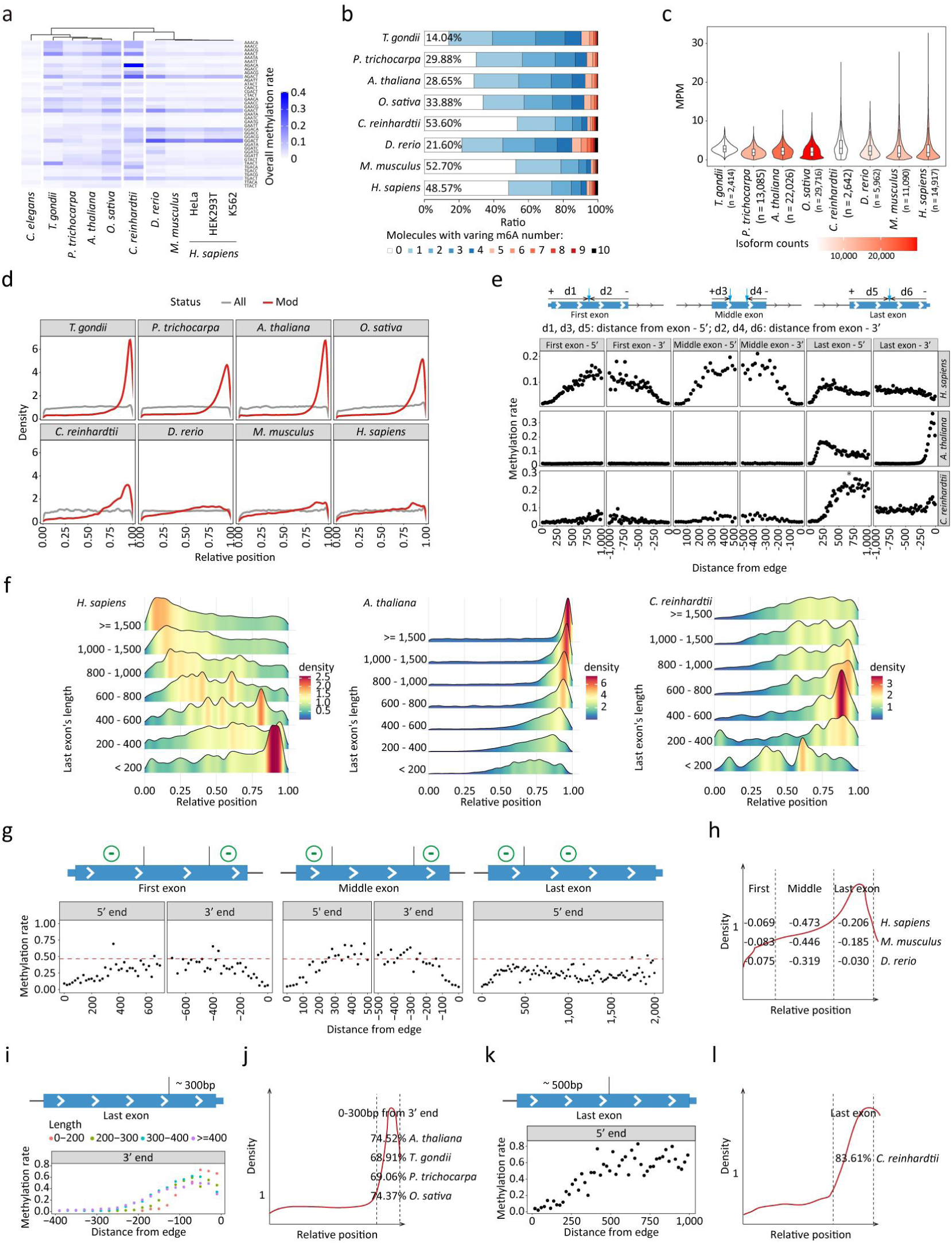
Quantitative comparison of single-molecule m6A profiles across multi-species. **a** Overall methylation rate of distinct motifs across various species. Clustering is based on “Pearson” distance. **b** Ratio of molecules with varying number of m6A in various species, and the ratios for molecules without m6A modifications (white bar) were also indicated. **c** Distribution of the MPM for different isoforms in various species. **d** Distribution of relative position of background (All) and modified (Mod) A bases within their respective molecules. **e** Overall methylation rate at positions with varying distance from one exon edge. **f** The distribution of relative position of m6A in last exons with varying lengths. **g** Top: schematic representation of exclusion deposition model in HEK293T. Bottom: m6A pattern shaped by exclusion deposition model and comparing the overall methylation rate at positions with varying distances from one exon edge to hypothetical default efficiency (red dashed lines), using GGACA motif as a representative. For first and middle exons, only potential motifs with distance from opposite edge greater than setting were retained for calculations, eliminating the influence of opposite edge. **h** Exclusion effects (the proportion of m6A content reduction to hypothetical m6A content) contributed by different types of exons in different vertebrates. **i** Top: schematic representation of the defined m6A deposition region in *A. thaliana*. Bottom: its m6A pattern and overall methylation rate at positions with varying distances from 3’ terminal, using AAACT motif as a representative. **j** The proportion of m6A within defined region in plants and protozoa. **k** Top: schematic representation of combination of defined m6A deposition and suppression region in *C. reinhardtii.* Bottom: its m6A pattern and overall methylation rate at positions with varying distances from 3’ ss of last exon, using AGACA motif as a representative. **l** The proportion of m6A in the last exon in *C. reinhardtii*.

Interestingly, the relative positions of m6A modifications on single RNA molecules display a preference towards the 3’ end across all species analyzed (Fig. 6d). The last exons contain around 40% of potential motifs but account for a significantly higher proportion of m6A modifications, ranging from 60% to 85% across species (Extended Data Fig. 6c). Consistently, the overall methylation rates and the proportion of last exons with at least one m6A site are much higher compared to other exons (Extended Data Fig. 6d-e). Nevertheless, the extent and region of this m6A enrichment at the 3’ end vary among different species (Extended Data Fig. 6f and Supplementary Fig. 12d). For instance, in plants and protozoa, m6A is sharply enriched in a small region near the RNA’s terminal, while in green algae, the enrichment region is larger and displays a higher level compared to vertebrates.

In line with recent discoveries^10–13^, we found that m6A deposition is suppressed within 150 bp from the splicing junction in human. This finding prompted us to conduct a comprehensive investigation into m6A modification at positions with various distances from the exon edges across species. Remarkably, we observed three categories of distinct m6A distribution patterns, which aligns with the clustering results based on motif preferences (Fig. 6e and Extended Data Fig. 7a-b). The first category, observed in vertebrates, is characterized by minimal m6A near the edges of both first and middle exons, with m6A levels remaining high in other regions of these exons (Fig. 6e and Extended Data Fig. 7a-b). However, due to their short lengths, most motifs are situated near the edges of these exons, resulting in relatively fewer m6A modifications. In contrast, the extended length of last exons allows them to accumulate more m6A modifications, although m6A levels are also low near their 5’ edge and gradually decrease towards the 3’ edge (Fig. 6e and Extended Data Fig. 7a-b). In the second category, observed in plants and protozoa, m6A is almost absent throughout the entire first and middle exons but confined to a small region at the 3’ end of the last exon (Fig. 6e and Extended Data Fig. 7a-b). In the third category, observed in green algae, m6A is confined to the entire last exon, while undergoes suppression within the first several hundred bp (Fig. 6e and Extended Data Fig. 7a-b). Comparing the distribution of m6A in last exons of varying lengths further supports the existence of these three distinct patterns (Fig. 6f and Extended Data Fig. 6g, 8).

### Quantitative elucidation of m6A distribution patterns

The recently proposed exclusion deposition model emphasized the key function of EJC in mediating m6A deposition^10–13^. Leveraging the single-molecule resolution of DRS, we aim to conduct a more rigorous and quantitative validation of this model and investigate whether it serves as the prevailing mode of m6A deposition in vertebrates. This model assumes that m6A is deposited with a default efficiency across transcripts, except in regions where EJC are present. To test this model, we first calculated the default deposition efficiency for various motifs. These values were inferred from the overall methylation rates in the central regions of middle exons, and were then compared with methylation rates at other positions (Fig. 6g, Extended Data Fig. 9a-b, 10a and Supplementary Fig. 13). Our results strongly support the EJC-mediated exclusion effect, which is expected to affect the middle exons, the 3’ end of first exons, and the 5’ end of last exons. In addition, we identified potential EJC-irrelevant exclusion effects in other regions. Specifically, m6A is also suppressed within a broader span of 400 bp at the 5’ end of first exons, and the entire last exon is subjected to additional suppression, which becomes more pronounced towards the 3’ end (Extended Data Fig. 9f, 9i and Supplementary Fig. 14). These findings extend the exclusion deposition model to the entire RNA molecule in vertebrates, which suggests other potential exclusion factors.

To quantify the contribution of the exclusion effect, we compared the observed m6A content with a hypothetical model where m6A is added with default efficiency across all regions. This analysis revealed that most motifs within the first and middle exons are located near exon edges and thus undergo strong suppression, resulting in an 80% reduction of m6A content in these exons (Extended Data Fig. 9c-h). In contrast, due to their longer lengths and moderate 3’ end suppression effect, last exons retain a higher m6A content (Extended Data Fig. 9i-k). Overall, approximately 75% of m6A content is reduced across the entire molecule in human and mouse, primarily due to the dense exon junctions in the middle exons and other potential exclusion factors in the last exons. The magnitude of these exclusion effects varies across vertebrate species, with zebrafish exhibiting a less pronounced reduction in m6A content compared to human and mouse (Fig. 6h).

Beyond vertebrates, we identified a completely different pattern in plants and protozoa, where m6A modifications occur explicitly within specific regions of the last exons. We further defined this region as spanning approximately 300 bp from the RNA terminal (Fig. 6i, Extended Data Fig. 10b and Supplementary Fig. 15). Remarkably, this small region accounts for 69%-75% of the total m6A content in these species (Fig. 6j and Extended Data Fig. 10c). In green algae, we also identified a distinct pattern, where after an initial 500 bp of suppression region, m6A is evenly distributed in the last exon, accounting for 83.6% of the total m6A content (Fig. 6k-l, Extended Data Fig. 10d and Supplementary Fig. 16). These distinct patterns observed in plants, protozoa, and green algae cannot be explained solely by the exclusion deposition model proposed in vertebrates and warrant further investigation to elucidate the underlying mechanisms.

## Discussion

As a modification on RNA, m6A introduces a remarkable level of complexity to the cellular transcriptome. In the past decade, significant advancements in m6A research have unraveled insights into its genomics, functions, and regulatory mechanisms^5^. Concurrently, increasingly sophisticated m6A mapping methods utilizing NGS platforms have emerged, advancing from peak-level to single-nucleotide resolution and from qualitative measurement to absolute quantification, with the latest GLORI method marking a milestone. However, as RNA exists in multiple copies within cells, the intrinsic heterogeneity among RNA molecules is challenging to completely capture using short-read sequencing. In this context, we introduce SingleMod, a cutting-edge approach that achieves the highest resolution and faithfully depicts m6A modifications on individual RNA molecules. We have harnessed SingleMod to acquire single-molecule m6A profiles across multiple species with distant phylogenetic relationship. We anticipate that, with the growing prevalence of the TGS platform^36^, researchers will easily obtain single-base and single-molecule resolution m6A maps in various research scenarios involving different species and different biological processes. Through DRS technology and corresponding computational tools, we are presented with a unique opportunity to obtain a holistic view of the transcriptome, including dimensions such as RNA splicing, polyadenylation, modifications, and more, that collectively contribute to the emerging field of epitranscriptomics.

Given the scarcity of fully-methylated sites in real samples^14^, we developed a multi-instance regression framework to train SingleMod. This framework effectively leverages information from sites exhibiting low to high m6A levels, facilitating comprehensive model training. Considering that most known RNA modifications, such as pseudouridine and m5C, exhibit low stoichiometry, we believe that this innovative framework could be particularly valuable for studying other RNA modifications. In practice, the specificity of m6A and the limited availability of data restricted our training of SingleMod to 39 prominent motifs, although we have integrated data from multiple species. While m6A in most known species explicitly occurs within DRACN motifs^6^, applying our model to a species without prior knowledge may introduce bias, particularly in terms of quantitative assessment. Similarly, given the current false positive rate and limited motif scope, identifying *METTL3*-independent m6A sites, which are typically extremely rare, remains a significant challenge for DRS-based tools. Moreover, inherent limitations of DRS, such as truncated reads and irregular current signals at the end of molecules, would hinder the accurate identification of RNA modifications in these regions. We anticipate that future research focusing on accumulating more comprehensive data across a wider range of species and exploring innovative synthetic data generation methods could help overcome these limitations. Despite these ongoing challenges, SingleMod represents a significant advance in the field, achieving unprecedented single-molecule resolution and contributing to more accurate and comprehensive m6A maps compared to existing methods.

Our ability to quantify m6A levels at single-molecule resolution has led to a reevaluation of previous quantitative findings based on traditional methods^30^. Because traditional methods average modifications across a pool of transcripts, they may overestimate the true m6A content on individual RNA molecules. Strikingly, approximately 50% of RNA molecules in our study exhibit no m6A modifications. In general, single-molecule m6A quantification is lower than previously estimates provided by traditional methods, such as LC-MS/MS. However, we should note that those methods cannot accurately determine the precise number of m6A modifications on individual RNA molecules. Our study also highlighted significant m6A heterogeneity among RNA molecules from a given gene, with m6A modifications tending to be distributed across multiple molecules and sites. Despite this heterogeneity, our research indicates that m6A promotes RNA degradation or regulates pre-RNA processing irrespective of its specific locations, functioning in an additive manner. Confirming this hypothesis in future research will be crucial for fully understanding the mechanisms by which m6A exerts its functions and for identifying potential target sites for functional disruption^37^. This inherent heterogeneity also provides a unique opportunity to study the interaction between m6A and pre-RNA processing, or other aspects of gene regulation. By categorizing molecules into different groups based on their m6A signatures and splicing (or PAS) structures, we can directly compare these naturally occurring “subpopulations” without the need for genetic manipulation. While we provided compelling molecule-level evidence for certain m6A and AS (or APA) interactions, it is essential to remain cautious regarding potential indirect regulatory effects.

Our comparative analysis of single-molecule m6A profiles across multiple species has revealed three distinct m6A distribution patterns. In vertebrates, we identified a combination of the well-established EJC-mediated exclusion and uncharted exclusion effects at the 5’ terminal and the last exon, contributing to a substantial 75% reduction in m6A content. While the exact mechanisms driving these additional exclusion effects remain to be fully elucidated, we speculate that certain RNA-binding proteins (RBPs) such as cap-binding proteins and transcription termination complexes may play roles in this process. In contrast to the exclusion effect observed in vertebrates, the distinct m6A patterns observed in plants and protozoa suggest a contrasting effect, through which the m6A writer may be recruited by unidentified factors, possibly including RBPs, to specifically add m6A at the RNA terminal. In green algae, the observed pattern suggests that certain 3’ UTR-preferred RBPs may be involved in directing the m6A writer to the last exons. Furthermore, m6A is also suppressed in a broad region at the beginning of the last exons, potentially through a mechanism akin to the EJC-mediated exclusion observed in vertebrates. Therefore, we speculate that m6A landscapes may be shaped by “inclusion” effects or the combination of both “inclusion” and “exclusion” effects in non-vertebrate species.

Overall, our study precisely delineates the m6A landscape at the highest single-base and single-molecule resolution across diverse species, offering valuable mechanistic insights for future research. The development of DRS technology and corresponding data analysis tools, such as SingleMod, has established a powerful new research paradigm for comprehensively analyzing the epitranscriptome.

## Methods

### Sample collection and RNA preparation

The mouse embryonic stem cells (mESC) were provided by our collaborator (Dr. J. Chen from the Guangzhou Institutes of Biomedicine and Health). A detailed description can be found in a previous study^38^. HEK293T cells (ATCC, passages <25) were cultured in DMEM (Corning; Cat#: 10-013-CVRC) supplemented with 10% fetal bovine serum (BIOVISION; Cat#: BVS500) at 37°C with 5% CO2. All cell lines used were confirmed the absence of mycoplasma contamination. The seeds of wild type *A. thaliana* accession Col-0 were sown and the seedlings were harvested in accordance with the experimental conditions outlined in a previous publication^28^. Rice (Oryza sativa L. subsp. japonica cultivar Nipponbare) was grown and roots from the two-week-old seedlings were collected in accordance with the experimental conditions outlined in a previous publication^39^. *C. reinhardtii* cells were collected by centrifugation and lysed in TRIzol reagent (Thermo Fisher Scientific, 15596018). After extracting total RNA from the samples following the manufacturer’s guidance, mRNA was isolated via a Dynabeads mRNA Purification Kit (Thermo Fisher Scientific, 61006) with genomic DNA removal using TurBO DNase kit (Thermo Fisher, AM2238).

### Data collection

The training and evaluation of SingleMod were conducted using nanopore DRS data and the corresponding GLORI data from multiple samples (HEK293T, HeLa, mESC, Arabidopsis seedlings, rice root, Chlamydomonas). eTAM-seq data from mESC and DRS data from three types of m6A-control RNA molecules (Curlcakes, Oligomer and SL-Oligomers) were also included for SingleMod evaluation. In addition, DRS data from 5 other species (and K562) were included in the comparative analysis of multi-species single-molecule m6A profiles. Some of the data were generated by this study, while others were obtained from publicly available datasets:

1. HEK293T. Triplicate DRS data of WT and *METTL3* KO cells were collected from Ploy N. Pratanwanich et al., 2021^40^ (ENA: PRJEB40872) and then merged respectively; GLORI data was from Cong Liu et al., 2022^14^ (GEO: GSE210563).
2. HeLa. Triplicate DRS data were collected from Ploy N. Pratanwanich et al., 2021^40^ (ENA: PRJEB40872); GLORI data was from Cong Liu et al., 2022^14^ (GEO: GSE210563).
3. *M. musculus*. DRS data of WT and *mettl3* KO mESC were collected from Zhendong Zhong et al., 2023^21^ (GEO: GSE195618); additional DRS data of WT was collected from Piroon Jenjaroenpun et al., 2020^41^ (GEO: SRP166020); two DRS data of WT were merged for SingleMod evaluation; GLORI data was generated by this study. eTAM-seq data was from Yu-Lan Xiao et al., 2023^18^ (GEO: GSE201064).
4. *A. thaliana*. Quadruplicate DRS data of WT (Col-0) and KO (mutant defective in the function of a conserved m6A writer complex component *VIRILIZER*) were collected from Matthew T Parker et al., 2020^28^ (ENA: PRJEB32782) and then merged respectively; GLORI data was generated by this study.
5. *O. sativa*. DRS data of the root were collected from Feng Yu et al., 2023^39^ (GSA: CRR521994) and also generate by this study, and were merged together; GLORI data was generated by this study.
6. *C. reinhardtii*. Paired DRS and GLORI data were generated by this study and used for model training. We also generated additional DRS data used for single-molecule m6A profile analyses.
7. K562. DRS data were collected from Ploy N. Pratanwanich et al., 2021^40^ (ENA: PRJEB40872).
8. *D. rerio*. DRS data of the embryo were collected from Oguzhan Begik et al., 2022^42^ (ENA: PRJEB53494).
9. *P. trichocarpa*. DRS data of stem were collected from Yubang Gao et al., 2021^24^ (GEO: SRR12822922).
10. *T. gondii*. DRS data were collected from Dayana C Farhat et al., 2021^43^ (NCBI’s SRA: PRJNA705300). Fast5 files were provided by Dr Christopher Swale.
11. *C. elegans*. DRS data of adult *C. elegans* were collected from Nathan P.Roach et al., 2021^44^ (ENA: PRJEB31791).
12. Curlcakes. DRS data were collected from Huanle Liu et al., 2019^23^ (GEO: GSE124309). Curlcakes are comprised of four synthetic sequences (Curlcake 1-4) that include all possible 5-mers. m6A-modified RNA molecules were synthesized via IVT by substituting ATP with m6ATP.
13. Oligomer. DRS data were collected from Rachael E. Workman et al., 2019^20^. Oligomer containing a m6A modification was chemically synthesized and then ligated to carrier molecule (firefly luciferase transcript).
14. SL-Oligomers. DRS data were collected from Adrian chan et al., 2024^29^ (ENA: PRJEB74106), and data corresponding to sequencing run ERR12770810 was used here. RNA oligos with a central m6A-modified or unmodified DRACH motif (AGACT or GGACA or GGACC) and flanking splint sequences are ligated to m6A-modified or unmodified heteropolymers by splint-assisted ligation, respectively.

The above DRS data were produced using RNA002 kit. We also generated DRS data (HEK293T and rice root) using the latest RNA004 kit for training SingleMod.

### Nanopore direct RNA sequencing and data processing

Nanopore direct RNA sequencing was conducted following instructions provided by Oxford Nanopore Technologies (Oxford, UK) using DRS kits (SQK-RNA002 or SQK-RNA004) and MinION or PromethION flowcells. For both the self-generated data and public data, raw signals were converted into fastq format through basecalling using Guppy (v6.3.7) with default parameters (using Dorado for RNA004 data). Reads that passed the quality threshold (7) and had a length greater than 25bp were then aligned to the reference genome (Supplementary Table 5) using Minimap2 (v2.22-r1101) with the parameter settings “-ax splice --secondary=no”. Samtools (v1.10) was used to filter supplementary alignments and convert sam files to the bam format. Nanopolish^45^ (v0.13.2) eventalign module was used to align raw signals to 5-mer bases with the parameter settings “--scale-events --samples --signal-index” (using f5c^46^ for RNA004 data). sam2tsv was used to extract bases quality and alignment results for each read from bam files. Raw current signals, current statistic values, and base quality were extracted for each motif within each read. These features were then used as SingleMod input for training or prediction. For SL-oligomers, we iteratively mapped the reference oligo sequences locally to each read using Biopython with penalty settings of ‘2, −1, 2, −1’ and a score threshold of 40. Subsequently, the mapped oligos were chained to infer the ligated sequence, and reads with distance between two adjacent hits of ≤7 bp and a length of ≥300 bp were retained for further analysis. BWA was used to map each read to its inferred reference, with key parameter settings of ‘-A2 -B1 −O2 -E1’.

### GLORI experiments and data processing

GLORI experiments were conducted following instructions provided in Cong Liu et al., 2022^14^, with an adjusted incubation temperature in deamination step of 25 °C. Subsequent to GLORI treatment, libraries were constructed using a one-pot RNA library preparation method. Briefly, 50 ng of GLORI-treated RNA sample was repaired using T4 Polynucleotide Kinase (NEB, #M0201S). Pre-adenylated 3’-adapter and 5’-adapter were sequentially ligated to the RNA using T4 RNA Ligase 2, truncated KQ (NEB, #M0373S) and T4 RNA Ligase 1 (NEB, #M0204S), respectively. cDNA synthesis was then performed using HiScript III 1^st^ Strand cDNA Synthesis Kit (Vazyme, #R312-01). Finally, the library was amplified using Q5 High-Fidelity DNA Polymerases with GC enhancer (NEB, #M0491S) and subjected to size selection using a 6% polyacrylamide gel. The resulting library was sequenced on the Illumina NGS platform, with PhiX included to balance base bias. Two biological replicates were used for each species. Raw reads were pre-processed using cutadapt (v4.1), seqkit (v0.8.1) rmdup module and fastp (v0.21.0) for adaptor trimming, deduplication and quality control as well as UMI trimming, respectively. Then, the clean data were processed using the RNA-m5C pipeline developed by Huang T et al., 2022^47^, with some modifications described below. The base conversion settings were A-to-G and T-to-C. We used Star (v020201) for alignment with key parameter settings “--outFilterMismatchNoverReadLmax 0.05 --outFilterMultimapNmax 20 --outFilterType BySJout --outFilterScoreMinOverLread 0.8”. We calculated the global A-to-G conversion rate from the mapping results and verified that all GLORI experiments achieved an A-to-G conversion rate ranging from 94% to 98% (Supplementary Table 4). All A sites with signal greater than 0.9 (representing the ratio of reads with no more than 3 non-converted A to all reads) and coverage greater than 15 after discarding non-converted reads were retained for methylation rates calculation. Subsequently, sites with a difference in methylation rate between two replicates not exceeding 0.05 were kept, and the average methylation rates of two replicates were used as the benchmarks for SingleMod training or evaluation.

### SingleMod

#### The framework of SingleMod

For a specific genomic location (A base), the m6A modification status for individual molecules is unknown. However, the methylation rate at that site, indicating the proportion of modified molecules, is available. For this type of data, we propose a multi-instance regression framework to train model that achieve single-molecule m6A prediction. For a specific motif, such as GGACT, the following conditions are known:

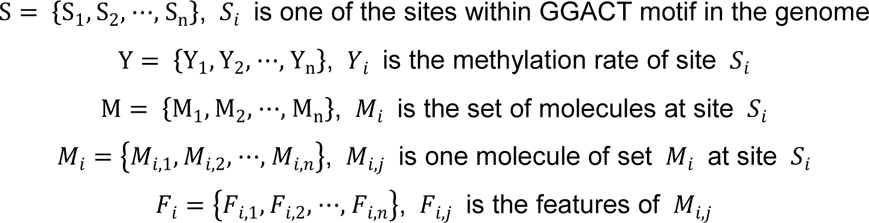

The modification status (*R_i,j_*) of molecule j at site i is unknown:

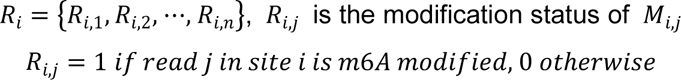

Given an initialized network N, the first step of our framework is to use network N to individually assess the modification status (*R_i,j_*) of each molecule *M_i,j_* of *M_i_* at site *S_i_*:

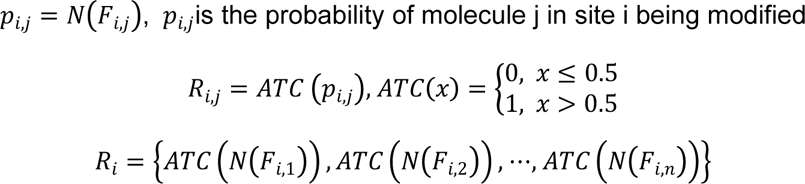

We obtain the prediction methylation rate of site *S_i_*:

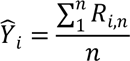

To optimize the parameters of the network, we minimize the mean squared error (MSE) between *Y_i_* and 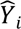 for all sites:

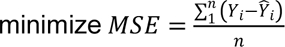

#### Informative features as input for SingleMod

For each molecule *M_i,J_* of *M_i_* at site *S_i_* its informative features are composed of five parts corresponding to the five positions of A/m6A reside in the pore during sequencing:

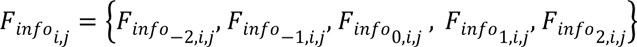

For example, assuming site *S_i_* is within the GGACT motif, the five positions of A/m6A reside in the pore can be represented as NNGGA, NGGAC, GGACT, GACTN, and ACTNN:

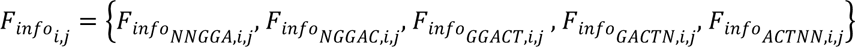

We also annotate features for each position with its 5-mer sequence information encoded through One-Hot encoding:

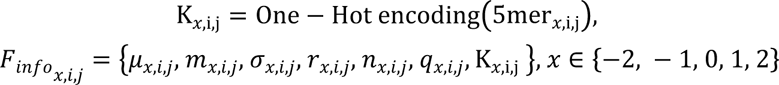

where 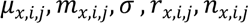 represent the mean, median, standard deviation, range and number of raw signals and q*_x,i,j_* represent the base quality from base-calling. Raw signals at each position can be extracted from the results file of Nanopolish eventalign. Typically, for each a position, it is measured multiple times when reside in the pore during sequencing, encompassing multiple raw signals, thus allowing for the calculation of the mean, median, standard deviation, range, and the number of these raw signals.

#### Raw signals as input for SingleMod

For each molecule M*_i,j_* of M*_i_* at site S*_i_*, input features are simple, consisting of a fixed-length segment of raw signals and corresponding sequence information. The center point of this fixed-length raw signals is the middle signal *R0,i,j* of the A/m6A to be detected. Then, the raw signals of a fixed length (e.g., 200) upstream and downstream of the middle signal can be extracted:

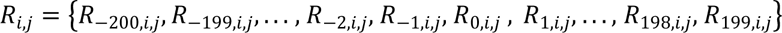

The sequence information (5-mer) corresponding to each raw signal is also encoded using One-Hot encoding:

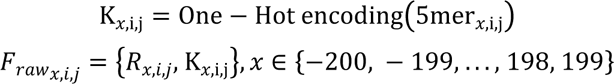

#### Determine the optimal input strategy and model architecture for SingleMod

We adopted a classification framework to explore the model inputs and architectures, operating in parallel for five m6A-rich motifs. Specifically, we utilized fully-methylated and unmethylated sites to generate positive and negative samples, respectively, which were then used to train a classification model after balancing the sample sizes. We employed the AUC of ROC to evaluate and compare the performance of various models. To begin with, we customized the model architecture separately for two input strategies: informative features and raw signals. We first determined the optimal combination of informative features through incremental feature selection. Meanwhile, we determined the optimal lengths of raw signals, which is 400. Based on this, we introduced additional networks, including Resnet, Remore-like network, GoogLeNet-like network, and compared the model performance across different input strategies and network configurations. We found that using raw signals directly as input yielded superior model performance compared to using informative features, and both our custom network and the GoogLeNet-like network performed the best. However, when we further compared these two networks by implementing them in the multiple instance regression framework, we observed that the GoogLeNet-like network converged very slowly. Even after training for 340 epochs, its performance was still not as good as our custom network trained for 170 epochs. Therefore, we decided to use raw signals as the model input and our custom network as the model architecture for SingleMod.

#### The model architecture of SingleMod

Raw signals and their corresponding One-Hot encoded sequences are separately fed into a five-layer one-dimensional convolutional neural network (CNN) with increasing numbers of channels (16, 32, 64, 128, 256) and a fixed kernel size of 3. Subsequently, a dropout rate of 0.5 is applied. The outputs are then merged and further processed by another five-layer one-dimensional CNN with a fixed number of 64 channels and progressively increasing kernel sizes (3, 3, 5, 7, 11). The resulting output is flattened, subjected to a dropout layer (0.5), and fed into a fully-connected layers (192, 2). The probabilities of m6A modification are obtained through the Softmax function and then transformed into binary values (0 or 1) using a self-activation function.

#### Training and testing

We trained models for different motifs separately, initially 13 motifs using merged data from HEK293T and HeLa, and finally 39 motifs using merged data from HEK293T, HeLa, Arabidopsis seedlings, rice root, Chlamydomonas. For each motif, we repeated the training process 3 or 4 times. Specifically, all sites were randomly divided into training, validation, and testing datasets in an 8:1:1 ratio (or 9:0.05:0.05 when using merged data from multiple species). During the training process, we set the batch size to 10 and employed the Adabound optimizer to minimize MSE between 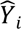 and Y_i_ for these 10 sites. We conducted training for 170 epochs and selected the model corresponding to the epoch with the minimum MSE on the validation set. We further assessed the model’s performance on the testing data using metrics including MSE, MAE, and Pearson’s r. The final models were selected based on their performance on independent evaluation data from mESC. It is important to note that only sites with a coverage of at least 20 DRS reads were included. Furthermore, NGS-based RNA-seq data were used to identify SNP and indels, and those sites near SNP or indels (±4 bp) were excluded. For RNA004 version SingleMod, we used data from HEK293T and rice root and trained models for 36 motifs.

### Compare SingleMod with existing tools

#### Apply SingleMod and existing tools to DRS data from mESC and Arabidopsis seedlings

After obtaining the SingleMod models trained on HEK293T and HeLa data, we evaluated their performance on DRS data from mESC and Arabidopsis seedlings and compared them to existing tools. Nanom6A^24^, Tombo^48^, m6Anet^27^, and xPore^40^ were used as described in Zhendong Zhong et al., 2023^21^. For m6Anet, we used the latest version, which capable of providing single-molecule level m6A predictions, and these results were used in our analyses. For DENA^25^, the “LSTM_extract.py” script was used to extract feature with key parameter “--windows 2 2”, and then “LSTM_predict.py” script was used to predict m6A methylation rates for all detected sites. For m6ABasecaller^49^, fast5 files were directly basecalled using the alternative m6A basecalling model, and the score indicating m6A modification of all A sites were extracted from basecalling results as described in the article. Absolutely quantitative m6A labels (methylation rates) from two orthogonal NGS-based methods, namely GLORI and eTAM-seq, were used as benchmarks.

#### Benchmark methylation rate predictions

We first compared the methylation rates calculated from single-molecule m6A predictions to NGS-based benchmarks. As some tools (SingleMod, DENA, Nanom6A, m6Anet) can only predict m6A modifications within certain motifs, we retained sites within 10 common motifs and with methylation rates exceeding 0.1, as determined by either NGS-based methods or DRS tools, for the computation of evaluation metrics, including MSE, MAE, and Pearson’s r.

#### Benchmark single-molecule m6A predictions

To evaluate single-molecule m6A predictions, we assigned the labels for molecules at fully-methylated (methylations rates > 0.99) and -unmethylated (methylations rates < 0.01) sites, as determined by GLORI or eTAM-seq, as methylated (1) and unmethylated (0), respectively. We extracted the probability (or score) of these m6A predictions from different tools to plot ROC and PR curves for the 10 common motifs. In the case of employing GLORI, we combined data from mESC and Arabidopsis seedlings for plotting. Note that, we downsampled unmethylated molecules to balance methylated and unmethylated molecules. We also employed two sets of DRS data from m6A-control RNA molecules to directly evaluate single-molecule m6A predictions. In these molecules, m6A (or A) are introduced to designed locations, which enable us to calculate the sensitivity (TPR) and specificity (1-FPR) of various tools. SingleMod, DENA, m6ABasecaller, m6Anet and Nanom6A were included in comparison as only these tools can provide single-molecule level m6A predictions.

#### Benchmark site-level m6A predictions

To assess site-level m6A predictions, genome-wide A sites covered by at least 20 DRS reads and NGS-based methods were kept to create a candidate list. Each tool or NGS-based method provided a methylation rate for each site in this list, classifying a site as m6A site if its methylation rate is above 0.1. If a tool does not report the methylation rate or the site fell beyond the tool’s motif scope, the site is labeled as non-m6A site by that tool. We used m6A sites identified by NGS-based methods to benchmark m6A-site predictions by various DRS tools. The F1 score is an evaluation metric that harmonizes precision and recall, offering a balanced assessment of the performance across all tools, and it is calculated using the following formula:

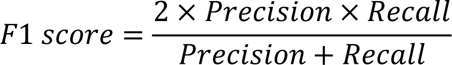

#### Compare SingleMod with Dorado

As Dorado only supports m6A calling from the latest 004 DRS data, we compared SingleMod with Dorado separately. We applied Dorado to HEK293T and rice DRS data for m6A calling using the ‘rna004_130bps_sup@v5.0.0’ configuration. Modkit was used to compute and extract methylation rates, which were then compared to GLORI benchmarks. Since the 004 version of SingleMod was trained on these data, for a fair comparison, we computed the evaluation metrics for SingleMod using its testing set, which was not involved in training.

### Single molecule m6A profile

#### m6A prediction at single-molecule level

The SingleMod predict module was used to predict m6A modifications within 39 motifs for each molecule, utilizing 39 individual SingleMod models corresponding to each motif. In-house Python scripts were employed to mark m6A modifications onto each read in the BAM file, allowing for visual inspection of single-molecule level m6A modifications in integrative genomics viewer (IGV).

#### Construct reference isoforms

Leveraging the advantage of long-read sequencing, our pipeline prioritizes the use of long reads to construct reference isoforms, with existing annotation file primarily used for correction and annotation. Specifically, bedtools (v2.26.0) was used to convert bam files to bed12 format. bedtools was used to merge 3’ end of each reads with key parameter “-d 5” and identified transcription termination regions. FLAIR^50^ (v1.6.1) correct module was used to corrects misaligned splice sites using genome annotations. Both corrected and inconsistent reads were retained and merged for subsequent use. FLAIR collapse module was used to generate intermediate files which were used to extract isoforms with key parameters settings “--gtf genome annotation --generate_map --keep_intermediate --check_splice --support 5 --trust_ends --isoformtss”. We extracted the non-single-exon isoforms already existing in gtf and isoforms with novel transcription termination sites (TTS) from the flair.collapse.firstpass.bed (using human as example: awk ‘$4 ∼ /ENST/ && $10 > 1’). Novel isoforms were extracted from flair.collapse.isoforms.bed (using human as example: awk ‘$4 !∼ /ENST/ && $10 > 1’). Single-exon isoforms with novel TTS were identified by comparing termination regions identified from reads with the 3’ end of single-exon isoforms in gtf. The above isoforms and the remaining isoforms in gtf were combined to construct the final reference annotations in bed12 format, with each isoforms annotated with its host gene.

#### Read to isoforms assignment

Trmap^51^ (v0.12.6) was used to found overlapped reference isoforms for each read, classifying their relationship based on splicing chain match type and calculated the exon overlap ratio. In-house Python script was used to assign each read to unique isoform and gene. Briefly, in the read-isoform relationship, if “=” (exact match of splicing chain) is present and the overlap ratio is greater than 0.7, the read is assigned to the corresponding isoform. If “=” is not present but “c” (contained in reference) exists and the overlap ratio is greater than 0.9, the read is assigned to the corresponding isoform. Lastly, if neither “=” nor “c” is present but “k” (containment of reference) exists and the overlap ratio is greater than 0.9, the read is also assigned to the corresponding isoform. If multiple instances of “=”, “c”, or “k” are present, the isoform with the highest overlap ratio is selected. For other cases, reads cannot be assigned to any isoform. However, we assigned these reads to the host gene corresponding to the isoform with the highest overlap ratio if overlap ratio was greater 0.9. Reads that could be successfully assigned to isoform or gene were considered as full-length reads and used for following analysis. As for gene expression quantification, reads were assigned to the host gene corresponding to the isoform with the highest overlap ratio.

#### m6A quantification

We quantified m6A modifications at different levels. At the cellular level, we analyzed the proportion of molecules with varying number of m6A modification. At the gene or isoform level, we calculated the proportion of modified molecules (those with at least one m6A modification, PMM), the average number of m6A modifications per molecule (MPM), and the average number of m6A modifications per molecule per 1,000 bp (MPKM). We calculated the counts per million (CPM) for each gene or isoform to represent their expression levels. It is important to note that CPM for direct RNA sequencing and transcripts per million (TPM) for short read sequencing serve as comparable metrics.

#### Sequence analysis of m6A

39 motifs from all molecules were counted and classified to “Unmodified” and “Modified” based on their modification status. Overall methylation rate of each motif was defined as the ratio of the number of “Modified” motifs to the total number of motifs. Seqlogo was used to plotted the composition of flanking sequence of m6A, which represented the motif enrichment at single-molecule level.

#### Relative position of m6A

This value was obtained by calculating the ratio of the distance from m6A position to the 5’ end of the molecule (or exon) to the length of the molecule (or exon). To define the region interval of start codon, last exon junction and stop codon, we applied a sliding window manner to determine the region interval when the quantity occupied half and the window size was minimized. For gene-specific m6A distribution, RNA molecules are divided into three equal intervals. For a specific gene, if over 25% of m6A modifications fall within one interval, it is deemed to have significant m6A modifications in that interval.

#### Overall methylation rate at positions with distance from landmarks and in different exon types

39 motifs from all molecules were counted and classified to “Unmodified” and “Modified” based on their modification status. For overall methylation rate at positions with varying distance to landmarks (start codon, last junction and stop codon), these values were derived by calculating the proportion of modified motifs that share the same distance (5 bp were grouped into a bin) from landmarks, relative to the total number of motifs. The overall methylation rate for different exon types, is calculated as the proportion of modified motifs within responding exon type.

#### Relationship of adjacent m6A

To assess interaction between adjacent m6A, m6A sites pair (both coverage >= 50 and methylation rate >= 0.1) with distance between them less than 100 bp were used. We compared the theoretically probabilistic ratio to the observed ratio of molecules with both sites being m6A modified to the total number of molecules. If they are significantly unequal, it indicates that the modification status of one site influences the status of adjacent site on a single molecule.

#### Assess m6A cluster at single-molecule level

m6A clusters (with at least two m6a sites) were identified by merging m6A sites (coverage >= 10, methylation rate >= 0.1 and modified molecules >= 3) using bedtools merge with key parameter settings “-d 100”. We then calculated the proportion of molecules with different number of m6A modifications within the cluster. We also counted the number of molecules containing m6A clusters with varying number of m6A modifications by merging adjacent m6A modifications with distance less than 100 bp within single RNA molecule.

#### RNA heterogeneity

To describe differences among molecules within a specific gene, we introduce the concept of heterogeneity. We quantify heterogeneity through the probabilistic sampling of two identical molecules within a gene, and a higher probability signifies reduced heterogeneity (heterogeneity = 1 - probability). We explore two facets of molecular heterogeneity: one arising from SS or PAS selections variations (isoform-related), and the other arising from m6A modifications.

#### Overall methylation rate at positions with varying distance from one exon edge

39 motifs from all molecules were counted and classified to “Unmodified” and “Modified” based on their modification status. These value were derived by calculating the proportion of modified motifs that share the same distance (20 bp were grouped into a bin) from exon edge. For the middle exons, when calculating the overall methylation rates toward one edge, only sites with a distance from the opposite edge greater than 200 bp were retained for calculation.

### m6A and RNA stability

RNA stability data were obtained from Qiushuang Wu et al., 2019^52^. To investigate the relationship between m6A and RNA stability, we simultaneously compared the RNA stability of genes with varying PMM, MPM, and MPKM. To investigate whether the role of m6A in promoting RNA degradation is influenced by its specific position, we compared RNA stability among genes with different primary m6A contribution intervals, while keeping MKPM constant. Specifically, we divided the molecules into three equal intervals and if a gene’s m6A content within one of these intervals exceeded 60%, it was considered that the primary source of m6A for that gene was that particular interval. When comparing m6A across different types of genes, we categorized protein-coding genes into regulatory genes and housekeeping genes, using the housekeeping genes set from the HRT Atlas v1.0 database^53^.

### m6A and RNA processing

#### Identify differential alternative splicing and alternative polyadenylation events upon *METTL3* KO

Utilizing the aforementioned outcomes of the “Read to isoforms assignment”, for each gene, we employed a modified version of call_diffsplice_events.py from FLAIR and in-house Python script es_detect.py to search for AS events and count reads with different splicing patterns for each of the three replicates in both WT and KO samples. As for APA events, we counted reads with different PAS selections according to transcription termination regions constructed in “Construct reference isoforms”. DRIMSeq^54^ (3.17) was used to calculate inclusion level (or PAS usage) and associated p-value. AS or APA events were considered differential between the WT and KO samples if the adjusted p-value < 0.05, and the change in inclusion level (max PAS usage change) >= 0.1.

#### Validate differential AS and APA events in *METTL3* inhibitor-treated HEK293T cells

HEK293T cells were cultured as mentioned above and then treated with either DMSO or 9 µg/mL STM2457 for 24 hours. After cell collection, total RNA was extracted with TRIzol reagent (Thermo Fisher Scientifific, 15596018) according to the manufacturer’s protocol. cDNA synthesis was performed using the HiScript III 1st Strand cDNA Synthesis Kit (Vazyme; Cat#: R312-01). qPCR was conducted using the HiScript II One Step qRT-PCR SYBR Green Kit (Vazyme; Cat#: Q221-01) on Kubo Tech q225 machine (Kubo; Cat#: q225-0268). We employed the methodology outlined by Peter Smibert et al., 2012^55^ for the validation of differential AS and APA events via RT-qPCR. Briefly, we designed two sets of qPCR primers: one set amplifies a region common to all isoforms in a given AS (or APA) event, and the other set amplifies the alternative region unique to some isoforms. We used the ΔΔCt method to quantify the relative inclusion level (or distal PAS usage) in AS (or APA) event between WT and *METTL3* inhibitor-treated cells, and the relative inclusion level (or distal PAS usage) of WT was set to 1. Two replicates were used in this experiment. All the primer sequences used in qPCR experiments are listed in Supplementary Table 7.

#### Multi-to-multi correlation analyses between m6A sites and AS or APA events

Within WT samples, there are naturally molecule groups with different modification status at specific sites and different SS or PAS selection in specific AS or APA events, allowing us to study the relationship between m6A and AS or APA events. Specifically, for each pair of m6A site and AS event (or APA), we performed Fisher’s exact test on four groups of molecules within the WT samples: 1) with m6A and the alternative region included; 2) with m6A and the alternative region excluded; 3) without m6A and the alternative region included; 4) without m6A and the alternative region excluded. If the p-value < 0.01, they were considered to be correlated.

### Quantification of exclusion effect

#### Validation of exclusion deposition model in vertebrates

We first calculated the overall methylation rate for each motif in regions that were not affected by EJC, specifically the central regions in middle exon with distance from both edges greater than 200 bp, to represent the default efficiency of m6A writer. Under the assumption that in exclusion deposition model, the methylation rates at regions where exclusion effect is absent would stabilize at the default efficiency of m6A writer, we then compared the overall methylation rate at positions with varying distance from one exon edge (20 bp were grouped into a bin) to the efficiency of m6A writer. When calculating the overall methylation rates toward the 5’ edge of first exons and both edges of middle exons, to eliminate the influence of the opposite edge, only motifs with distance from the opposite edge greater than 200 bp were retained. As for the 3’ edge of first exons, only motifs with distance from the opposite edge greater than 400 bp were retained.

#### Quantification of exclusion effect in vertebrates

We first calculated the exclusion effect for the first, middle, and last exons, respectively. Using middle exons and the GGACA motif as an example, we counted the number of GGACA motifs and the number of m6A-modified GGACA motifs (the observed m6A content) across all molecules. We then calculated the hypothetical number of m6A-modified GGACA motifs (the hypothetical m6A content) if without any exclusion effect: the number of GGACA motifs multiply by the default efficiency of GGACA motif. The reduction of m6A content by exclusion effect and the exclusion effect is calculate as follow:

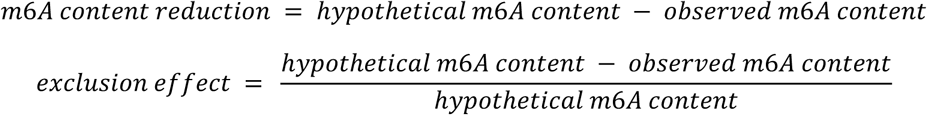

The overall exclusion effect in the middle exons is calculated by summing the observed m6A content and the hypothetical m6A content of each motif, and then applying the aforementioned formula. Likewise, the exclusion effect in the first and last exons is calculated using the same method. The total exclusion effect across entire molecule is calculated by summing the observed m6A content and the hypothetical m6A content of first, middle, and last exons, and then applying the aforementioned formula, and the contribution from different exon types is also calculated.

## Data availability

Sequencing data generated in this work can be found in GEO with the accession number GSE246632.

## Code availability

The source code and a detailed tutorial for SingleMod are publicly available at Github: https://github.com/xieyy46/SingleMod-v1.

## Acknowledgements

This work was supported by the Ministry of Science and Technology of China (2022YFC3400400, 2022YFA0912900 and 2019YFA0802203), National Natural Science Foundation of China (92253202, 32271499, 32270644, and 32100461), and Shenzhen Bay Scholars Program.

## Author Contributions

G.-Z.L. and Z.Z. conceived the project; Y.-Y.X. and Z.-D.Z. analyzed the data and wrote the manuscript; H.-X.C. conducted the experiments with the assistance from Y.-T.Q., Z.-H.R., Y.-L.L., F.W., B.-D.L., Y.S., D.Z., F.Y., Z.L. and J.W.; Y.-Y.X., Z.-D.Z., Z.Z., and G.-Z.L. revised the manuscript. All authors reviewed the results and approved the final version of the manuscript.

---

**Extended Data Fig. 1.**
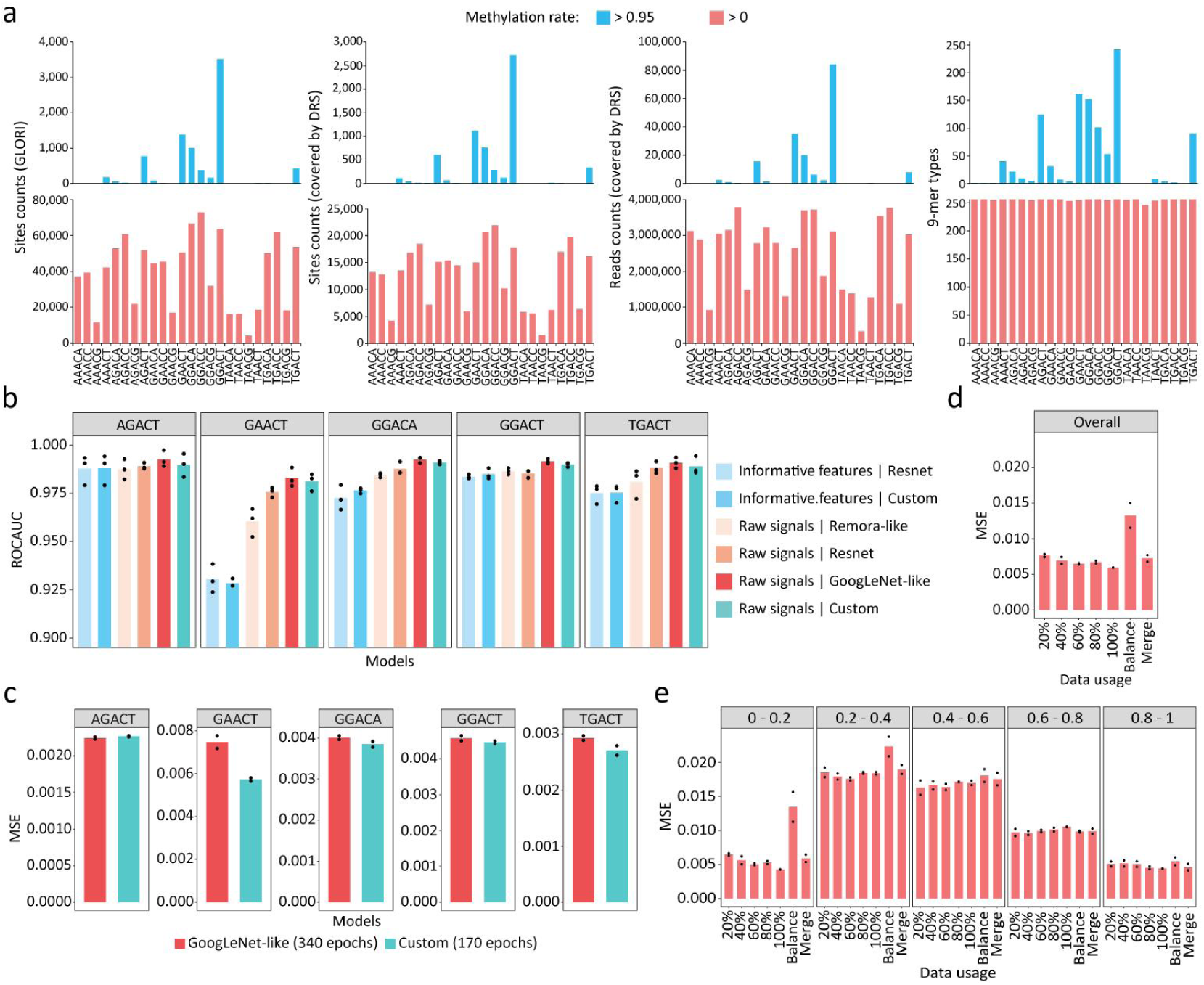
The development of SingleMod. **a** Statistics of GLORI and nanopore DRS data derived from HEK293T. From left to right: the number of m6A sites identified by GLORI; the number of these sites covered by DRS data; the number of DRS reads corresponding to these sites; the number of 9-mer types corresponding to these sites. **b** ROC AUC of models using different network architectures and input strategies across five motifs. **c** MSE of SingleMod using two best network architectures in **b** across five motifs. **d-e** Examining SingleMod’s overall performance (**d**) and its performance across various intervals (**e**), when incorporating varying proportion of low-methylation rate sites (<=0.05) for training.

**Extended Data Fig. 2.**
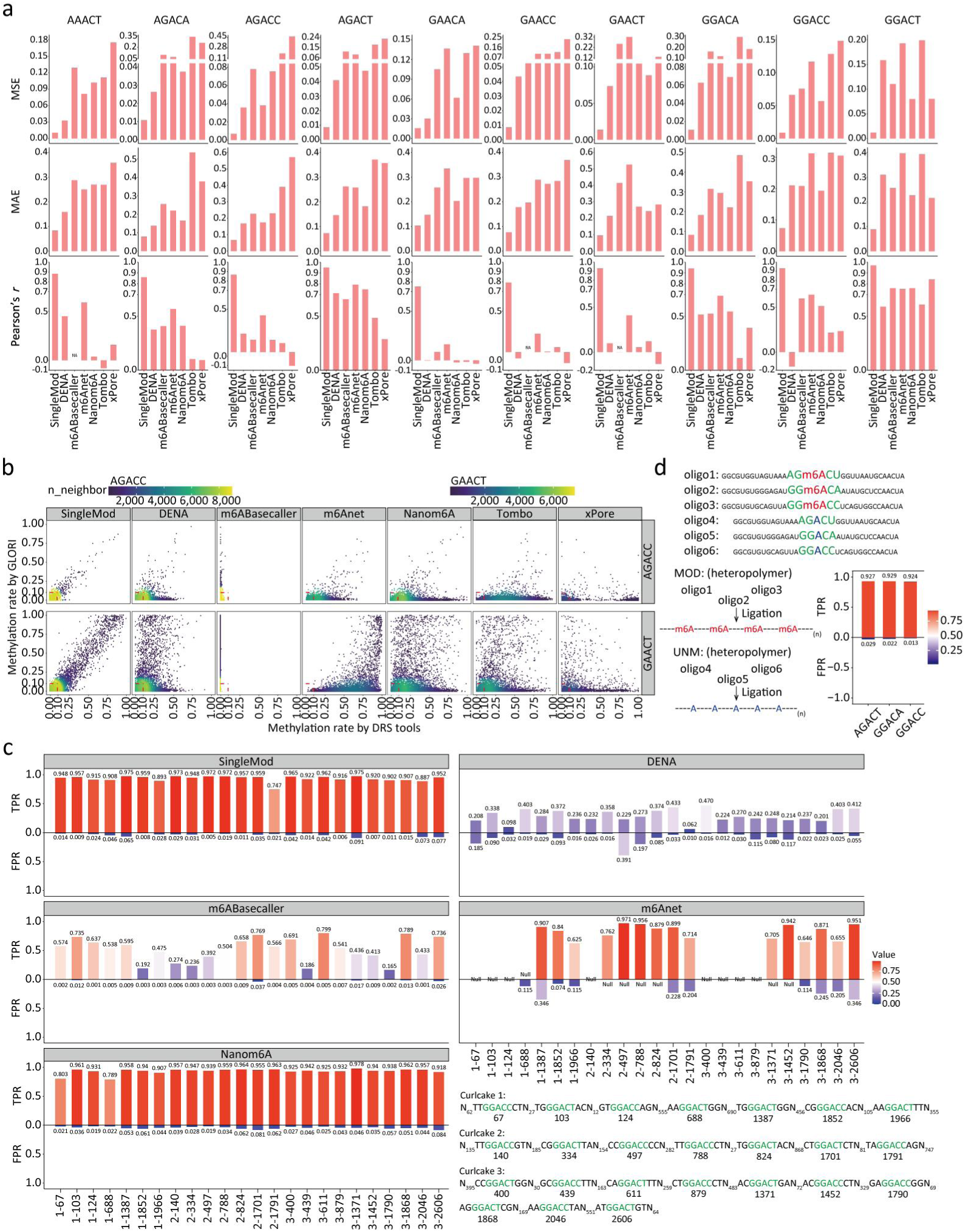
Comparison of the performance between SingleMod and other tools on different data. **a** The performance of SingleMod and other tools on various motifs. The Pearson’s r of m6ABasecaller in certain motifs was designated as “NA”, because this tool predicted these motifs in all molecules to be unmethylated. Data used in **a-b** were from *M. musculus*. **b** Comparing the methylation rates predicted by DRS tools to GLORI benchmarks. Two motifs AGACC and GAACT were used as representative. **c** Comparing the TPR and FPR of single-molecule level m6A predictions between SingleMod and other tools using IVT-derived (Curlcakes) m6A-control data. **d** TPR and FPR of single-molecule level m6A predictions by SingleMod using synthetic (SL-Oligomers) m6A-control data.

**Extended Data Fig. 3.**
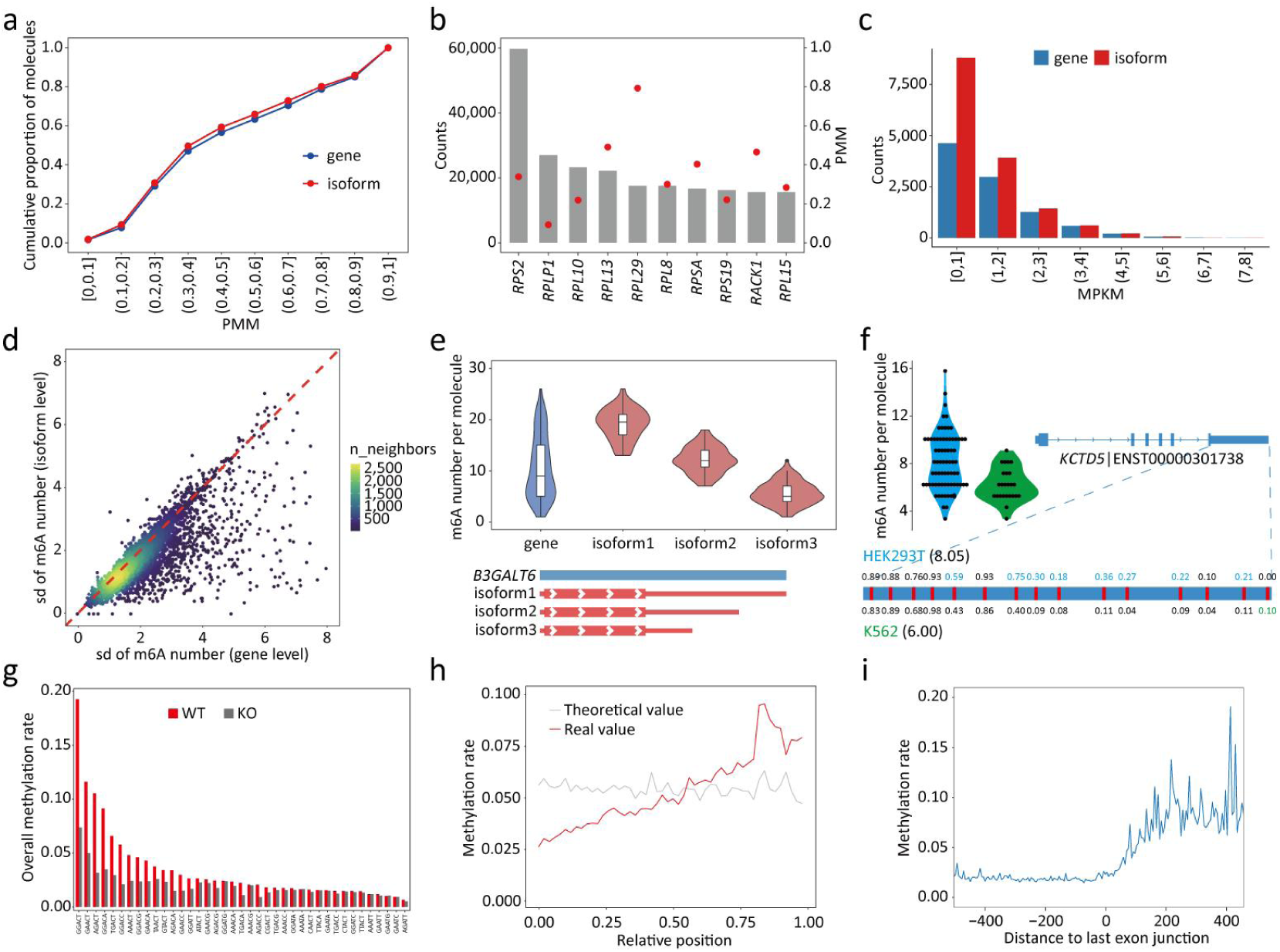
Single-molecule m6A profile in HEK293T. **a** Cumulative proportion of molecules from gene groups with varying PMM. **b** Molecules count (bar) and PMM (dot) for the top 10 highest expressed genes. **c** Counts of genes or isoforms with varying MPKM. **d** Comparing the standard deviation in m6A number among molecules between gene and isoform level (the mean standard deviation of multiple isoforms). **e** Distribution of m6A number in each single molecule from gene *B3GALT6* and its three isoforms. **f** Comparing the distribution of m6A number in all molecules of a representative isoform (left), the MPM of this isoform, and the methylation rates of each site (right) between HEK293T and K562. **g** Overall methylation rate of various motifs in WT and *METTL3* KO cells. **h-i** Overall methylation rate for DRACH motifs at different relative positions (h) and around last exon junction (i).

**Extended Data Fig. 4.**
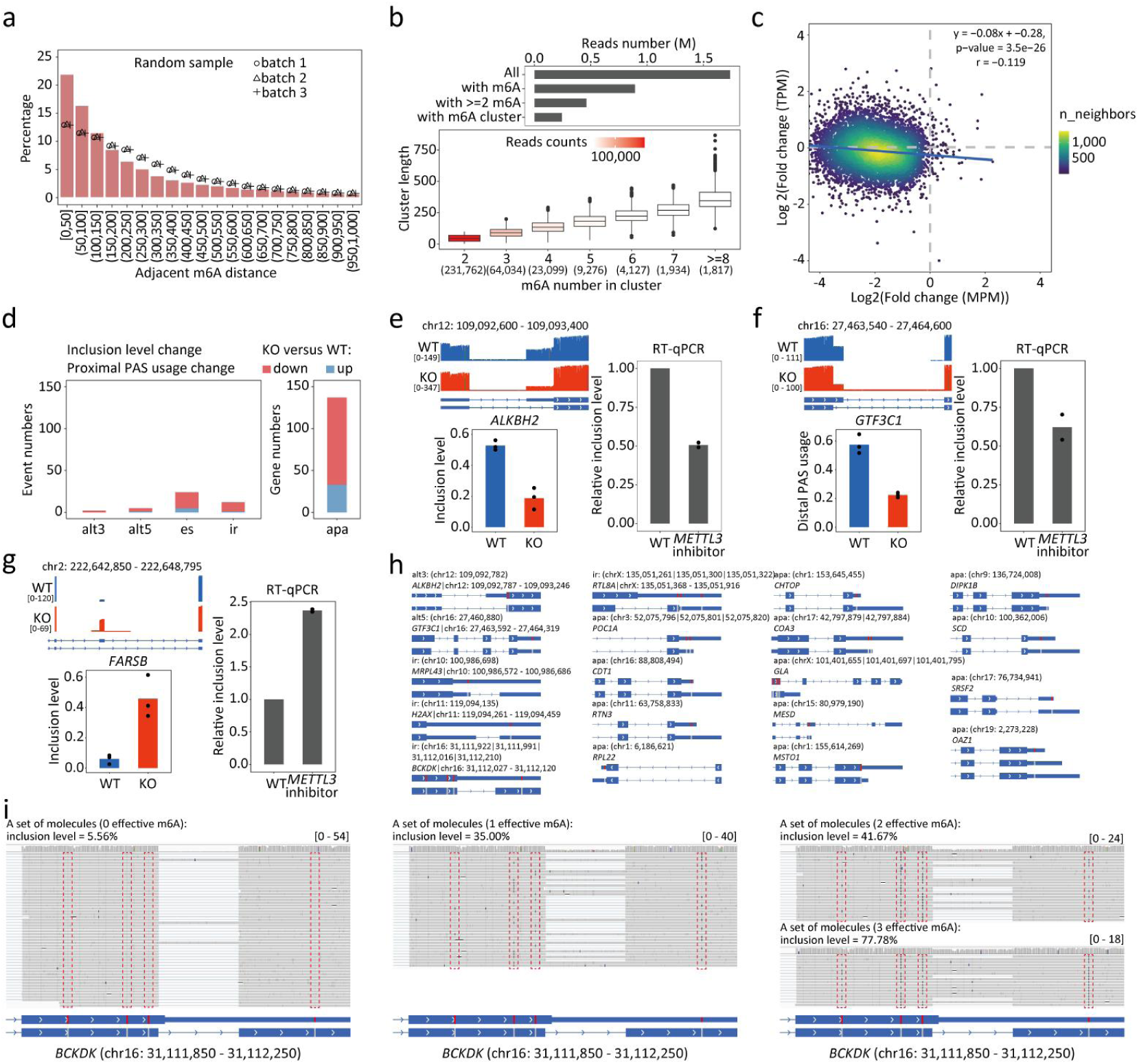
Differential AS and APA events upon *METTL3* KO and effective m6A sites in HEK293T. **a** Percentage of different distances between two adjacent m6A on individual RNA molecule. **b** Top: number of different types of molecules. Bottom: length of m6A clusters with varying m6A numbers and their corresponding molecules numbers. **c** Correlation between changes in gene expression and m6A level following *METTL3* KO. **d** Numbers of differential AS/APA events following *METTL3* KO. **e-g** Three examples of differential alt3 (**e**), alt5 (**f**) and es (**g**) events and their corresponding RT-qPCR validation results in cells treated with the *METTL3* inhibitor. **h** Differential AS/APA events and the specific locations of corresponding effective m6A sites identified through correlation analysis. **i** A representative case of **h**, showing a differential IR event in the gene POC1A and the effective m6A sites (red box). In the IGV snapshot which provided a visual representation that m6A affects splicing, molecules were grouped based on the number m6A modification at effective sites, and their corresponding inclusion levels were also indicated.

**Extended Data Fig. 5.**
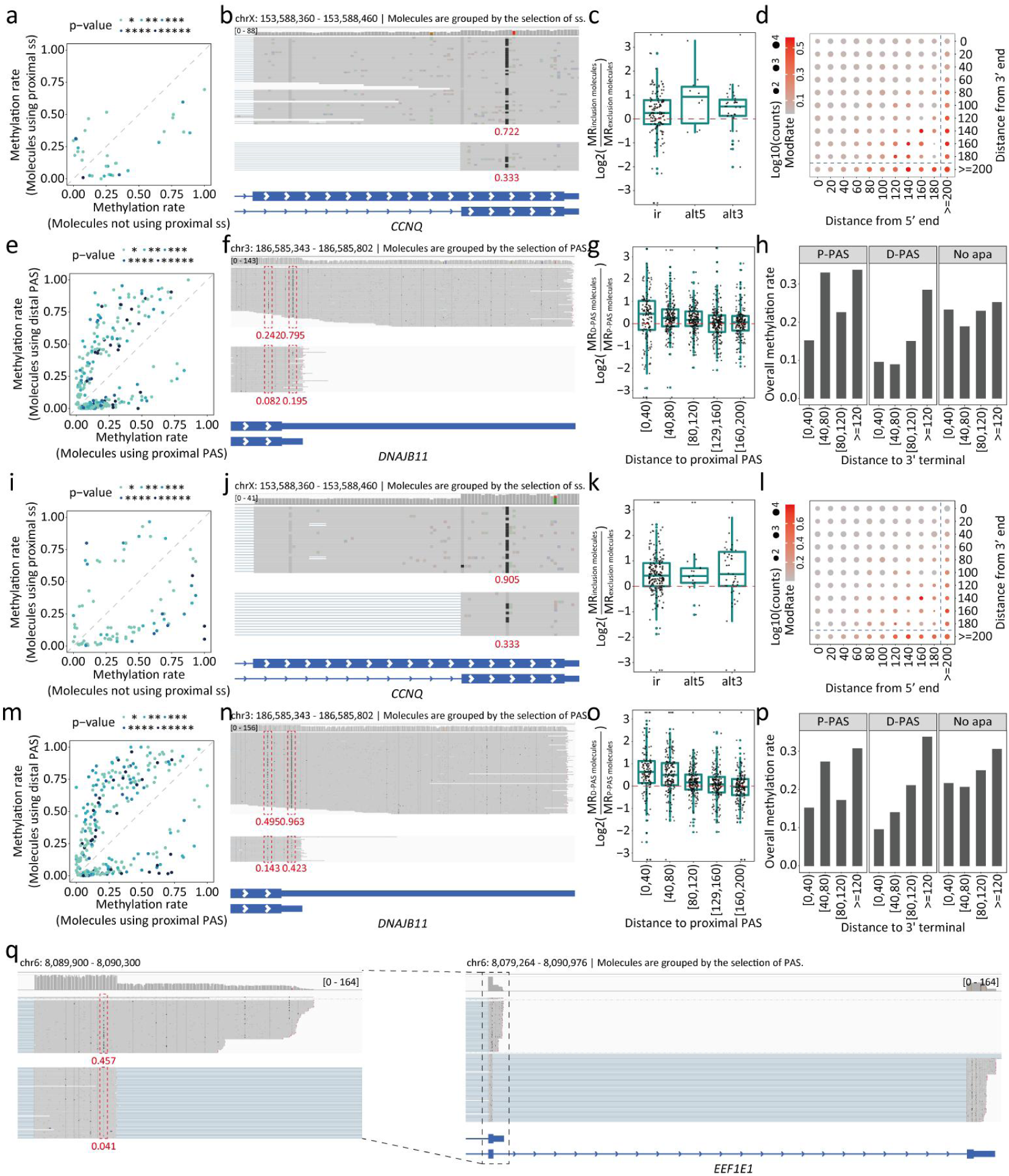
Single-molecule evidence that m6A deposition is affected by pre-RNA processing in human cell lines. **a** Dotplot compares the methylation rate of m6A site between two groups of molecules with different SS selections in the correlation analysis, as calculated in Fig. 5a. Each dot represents a significantly correlated m6A site-AS event pair, and its P-value was also indicated. Data used in **a-h** were from HeLa. **b** An IGV snapshot Illustrates a representative significantly correlated m6A site-AS event (alt3) pair. We grouped these molecules based on the 3’ SS selection and compared the methylation rates (indicated in red) of the correlated m6A site between two groups. **c** Comparing the methylation rates of nearby m6A sites (<=200 bp from exon edge) between inclusion-spliced molecules and exclusion-spliced molecules across different AS events, as showed in **Fig. a**. **d** Overall methylation rates at positions with varying distance from both edges in the middle exons, using GAACT motif as a representative. **e** Dotplot compares the methylation rate of m6A site between two groups of molecules with different PAS selection in the correlation analysis, as calculated in Fig. 5h. Each dot represents a significant correlated m6A site-APA event pair, and its P-value was also indicated. **f** An IGV snapshot Illustrates a representative significantly correlated m6A site-APA event pair. We grouped these molecules based on the PAS selection and compared the methylation rates (indicated in red) of the correlated m6A site between two groups. **g** Comparing the methylation rates of m6A sites near proximal PAS between D-PAS molecules and P-PAS molecules, as showed in Fig. 5h. **h** Overall methylation rate at positions with varying distance from the 3’ terminal of three types of molecules, using GAACT motif as a representative. **i-p** Results identical to those of **a-h**, but the data came from K562. **q** An IGV snapshot Illustrates a significantly correlated m6A site-APA event pair in gene EEF1E1, which is associated with intronic PAS and appears to be mediated by EJC. We grouped these molecules based on the PAS selection and compared the methylation rates (indicated in red) of the correlated m6A site between two groups.

**Extended Data Fig. 6.**
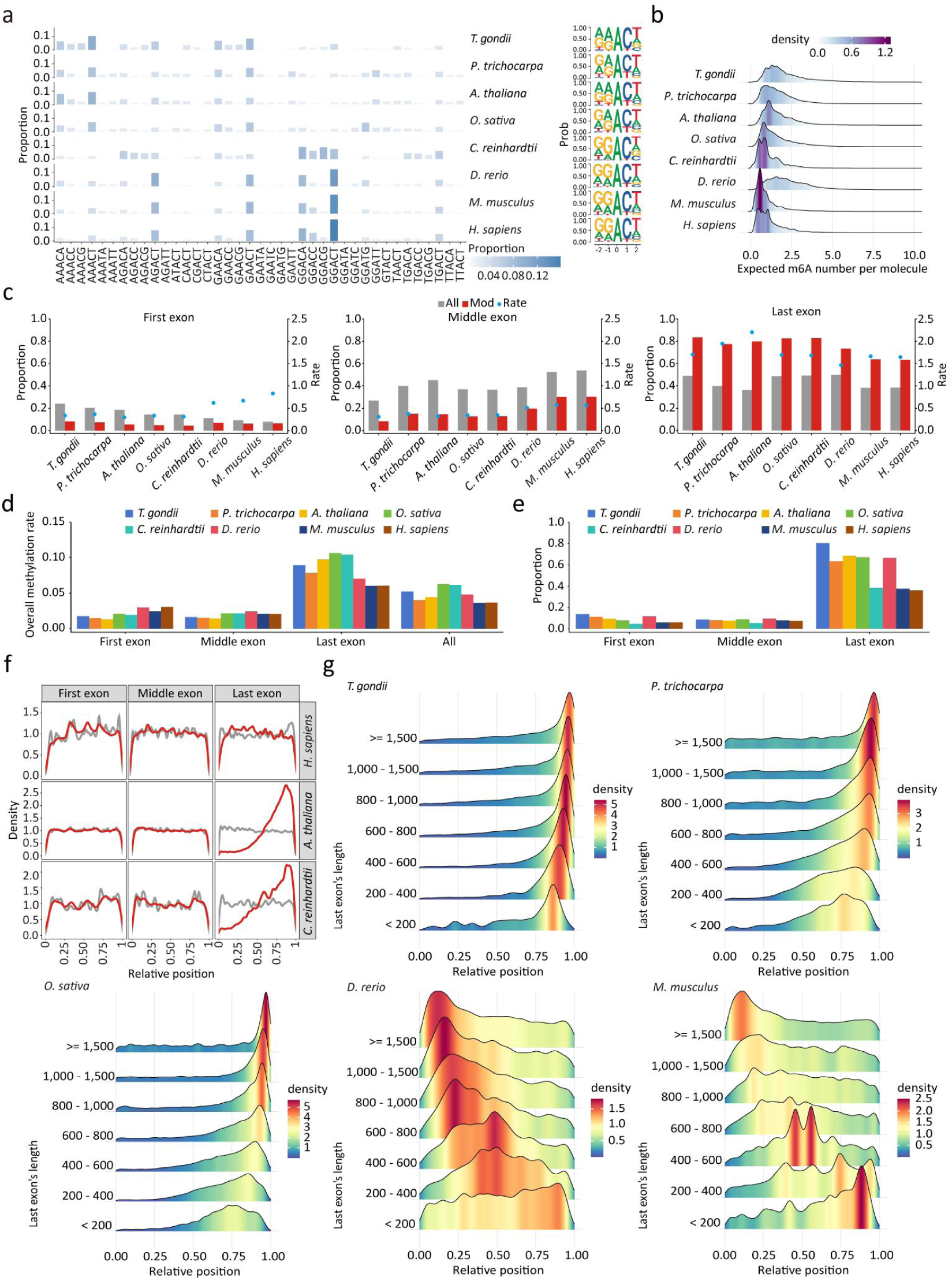
Single-molecule m6A profiles across multi-species. **a** Proportion of modified A bases within various motifs to all modified A bases. The composition of m6A flanking bases plotted by Seqlogo was shown on the right. **b** Distribution of the expected number of m6A modifications per molecule. The expected m6A number was calculated by summing up the overall methylation rates of each motif occurrence on a molecule. **c** Bar: proportions of background (All) and modified (Mod) A bases in distinct exons. Dot: the ratio between these two values. **d** Overall methylation rate in distinct exons. **e** Proportion of modified exons (containing at least one m6A). **f** The distribution of relative position of background (All) and modified (Mod) A bases within their respective molecules in different exons. **g** Distribution of relative position of m6A in the last exons with varying lengths.

**Extended Data Fig. 7.**
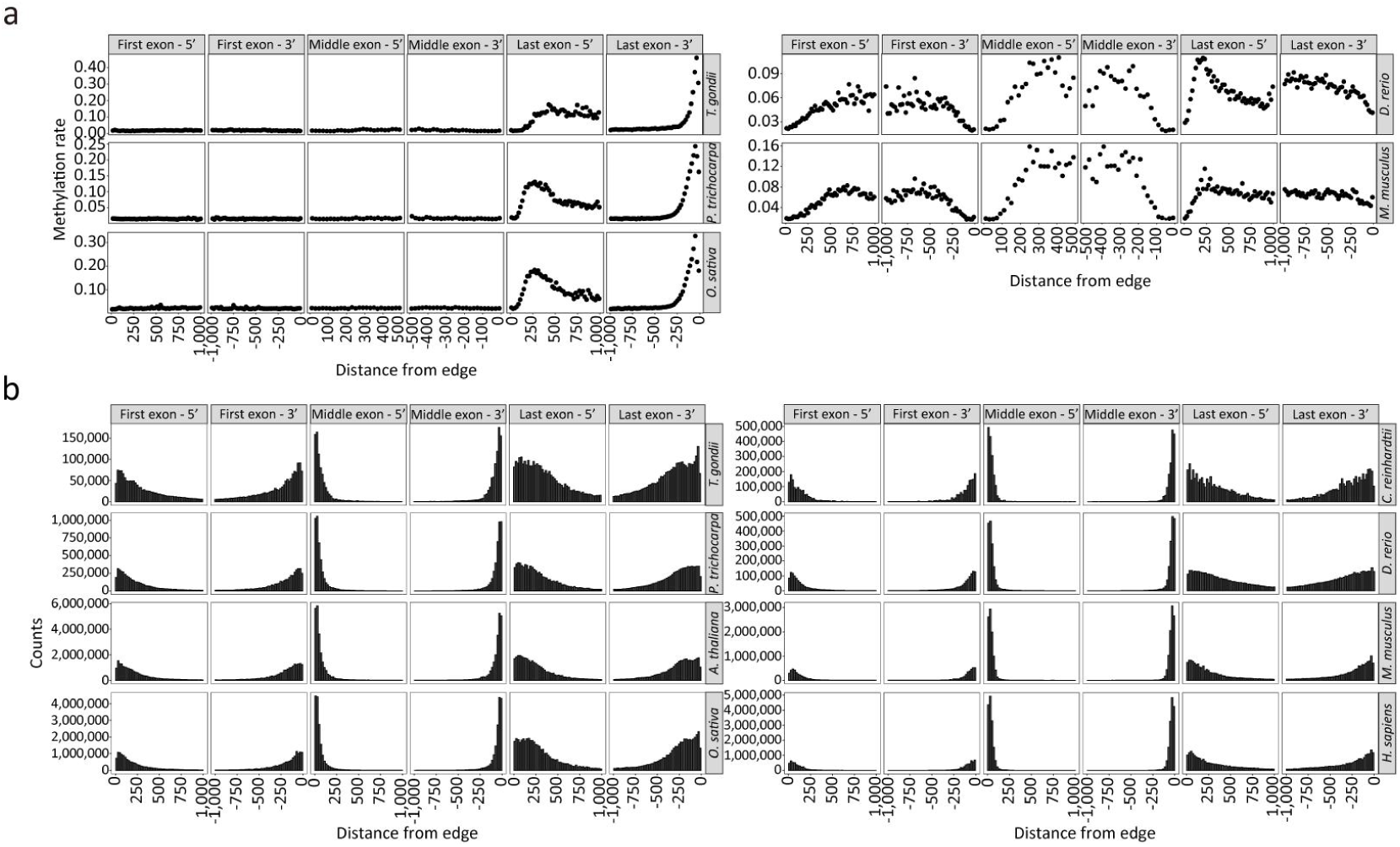
Overall methylation rate (a) and counts of motifs (b) with varying distance from one exon edge.

**Extended Data Fig. 8.**
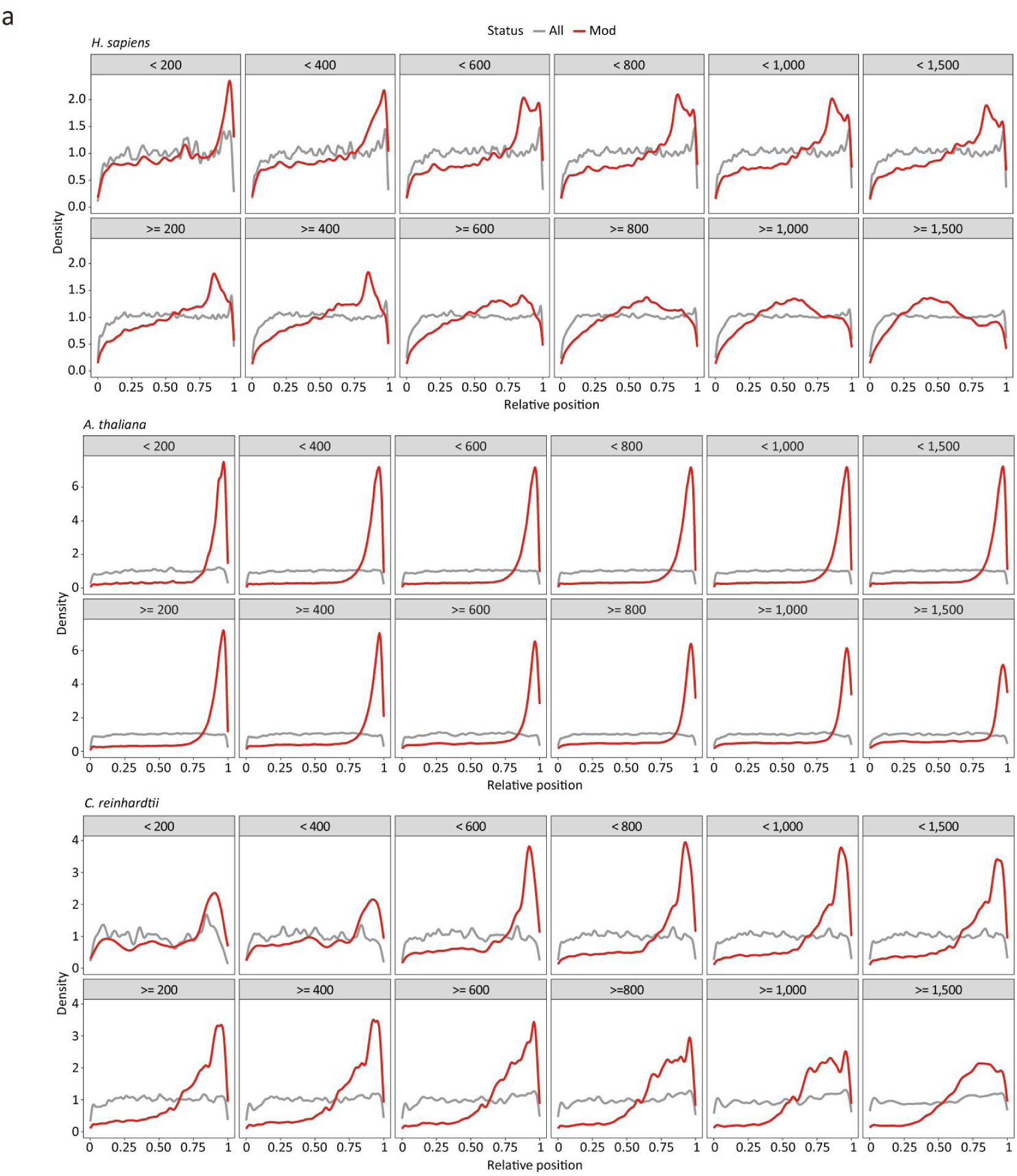
Distribution of relative position of background (All) and modified (Mod) A bases within their respective molecules with different last exon length.

**Extended Data Fig. 9.**
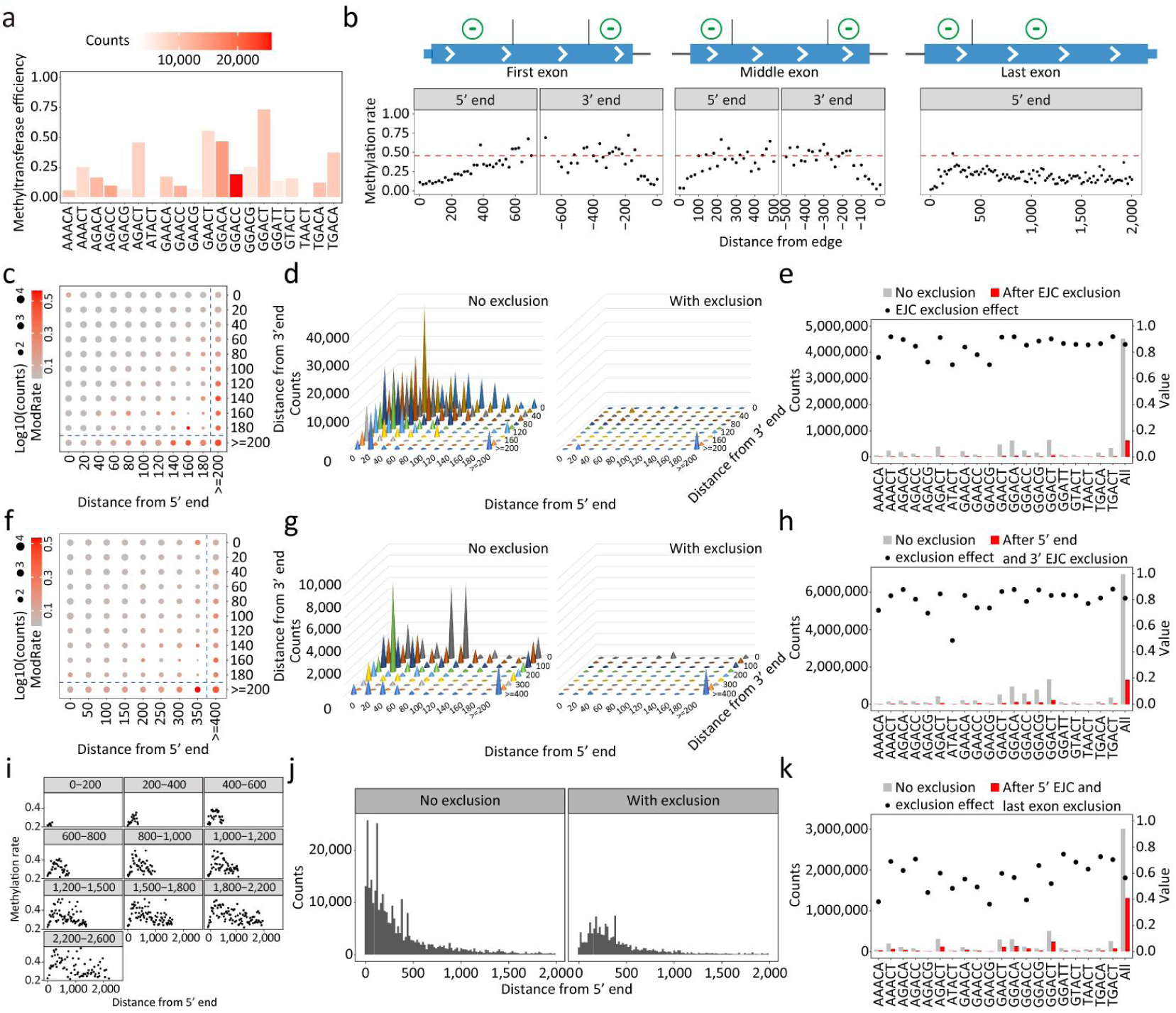
Quantitative analysis of exclusion deposition model in HEK293T. **a** Intrinsic efficiency of m6A deposition for top 20 motifs. **b** The same as Fig. 6a, but using AGACT motif as a representative. **c, f** Overall methylation rate and counts of motifs at positions with varying distance from both edges. **d, g** Hypothetical m6A content if without exclusion effect (left) and observed m6A content (right). **e, h** Bar: hypothetical m6A content and observed m6A content for various motifs. Dot: exclusion effect. **c-e** and **f-h** are results for middle and first exons, respectively, and **c, d, f, g** use GGACA motif as a representative. **i-j** Overall methylation rate at positions with varying distances from 3’ss in last exon with different lengths (**i**), and corresponding hypothetical m6A content (**j**, left) and observed m6A content (**j**, right), using GGACA motif as a representative. **k** Bar: hypothetical m6A content and observed m6A content in the last exon for all motifs. Dot: exclusion effect.

**Extended Data Fig. 10.**
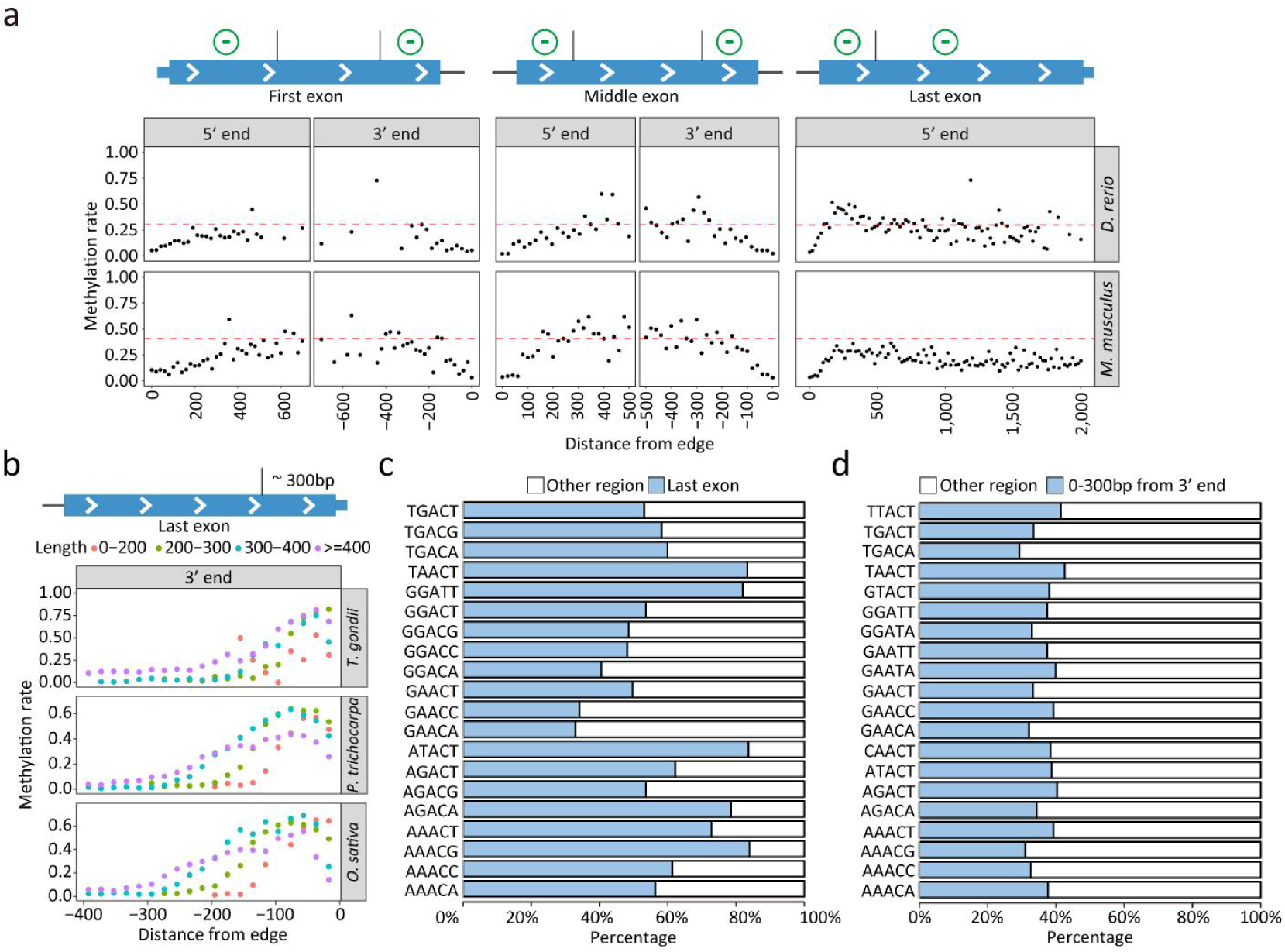
Quantitative elucidation of m6A distribution patterns across multi-species. **a** Top: schematic representation of exclusion deposition mode in vertebrates. Bottom: m6A pattern shaped by exclusion deposition model and comparing the overall methylation rate at positions with varying distances from one exon edge to hypothetical methylation rate (red dashed lines), using GGACA motif as a representative. **b** Top: schematic representation of the defined m6A deposition region in plants and protozoa. Bottom: their m6A patterns and overall methylation rate at positions with varying distances from 3’ terminal, using AAACT motif as a representative. **c** Percentage of motifs within and outside the defined m6A deposition region in *A. thaliana*. **d** Percentage of motifs within and outside the last exon in *C. reinhardtii*.

