## Supplemental Figures for "Single-Molecule Direct RNA Sequencing Reveals the Shaping of Epitranscriptome Across Multiple Species"

### Contents:

1 Supplementary Figures

2 Supplementary Tables

### SUPPLEMENTARY FIGURES

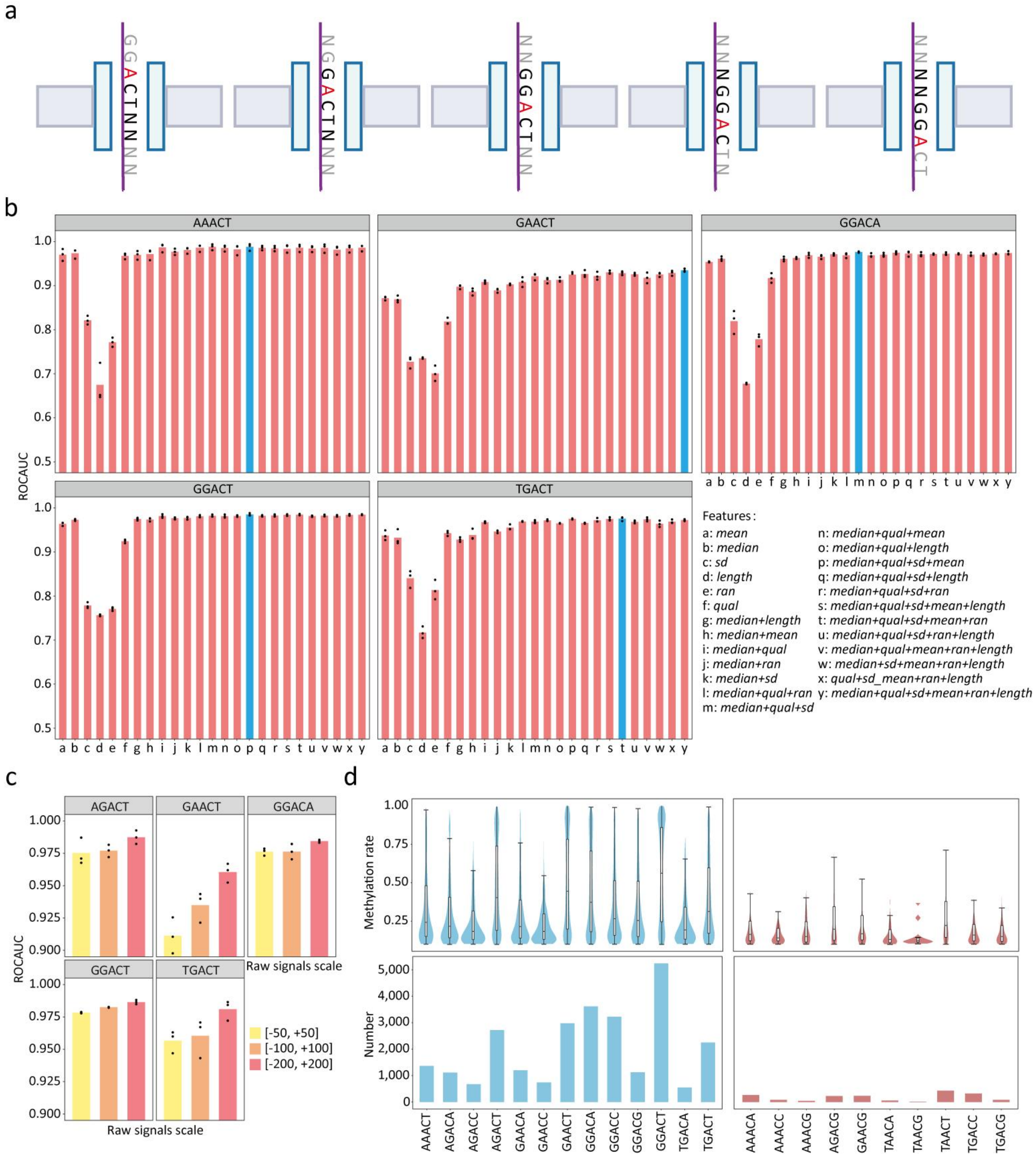


**Supplementary Fig. 1|Development of SingleMod. a** Schematic illustration depicting the five locations of A base and corresponding 5mer in the nanopore during sequencing. **b** ROCAUC of classification models using different combinations of features across 5 motifs. **c** ROCAUC of classification models using raw signals within different length scales across 5 motifs. **d** The distribution of methylation rates (top) and number (bottom) of m6A sites with methylation rate exceeding 0.1 in 23 DRACN motifs in HEK293T. Data was derived from GLORI results. Initially, SingleMod was separately trained using data from 13 predominant motifs (left).

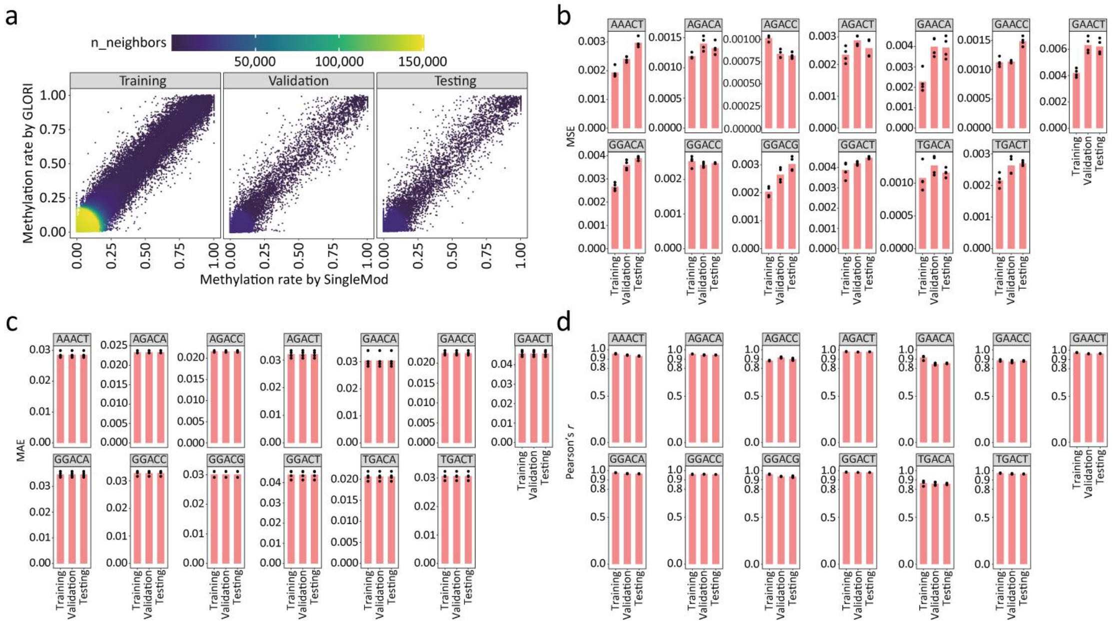

**Supplementary Fig. 2|Performance of SingleMod on different datasets from HEK293T. a** Comparing the methylation rates predicted by SingleMod to GLORI benchmarks. **b-d** The performance of SingleMod on various motifs in different datasets.

a

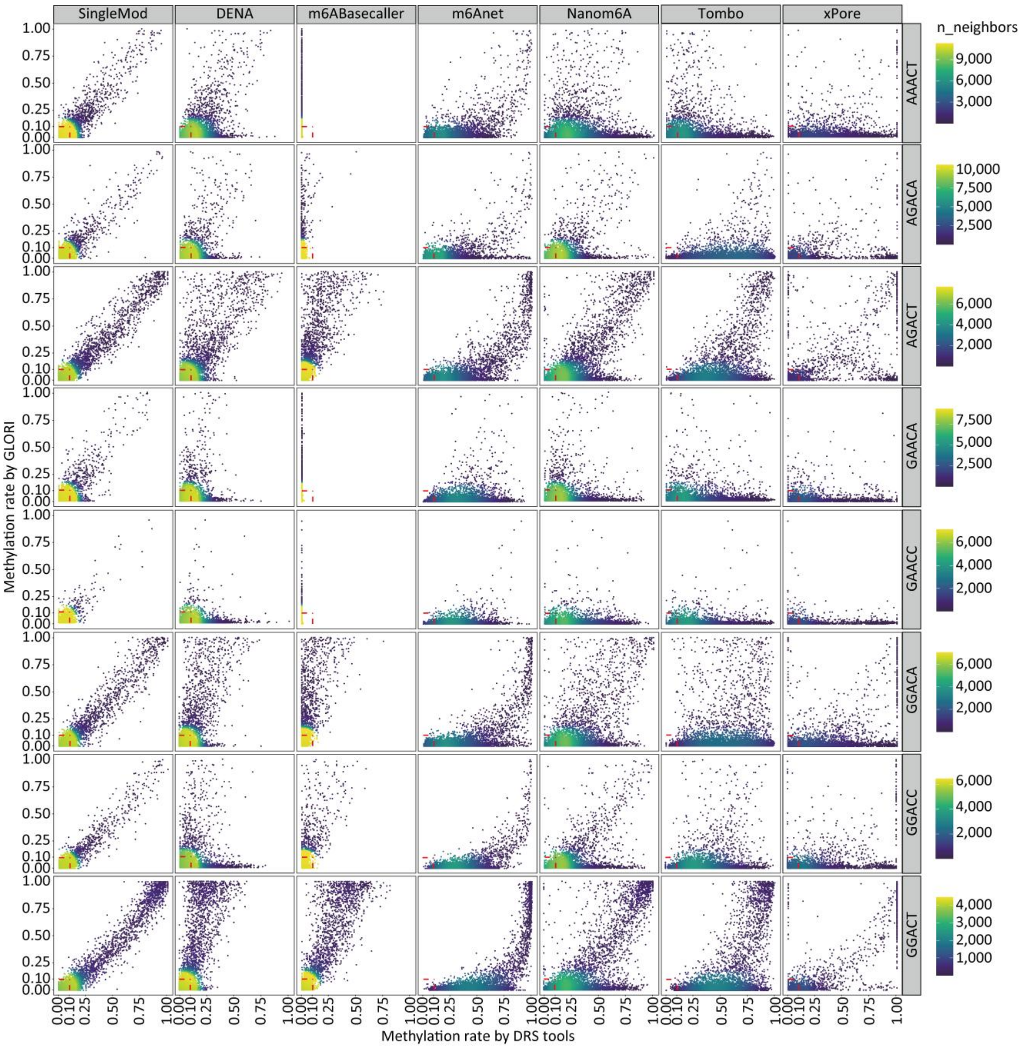


**Supplementary Fig. 3|Comparing the methylation rates in various motifs predicted by DRS tools to GLORI benchmarks on *M. musculus* data.**

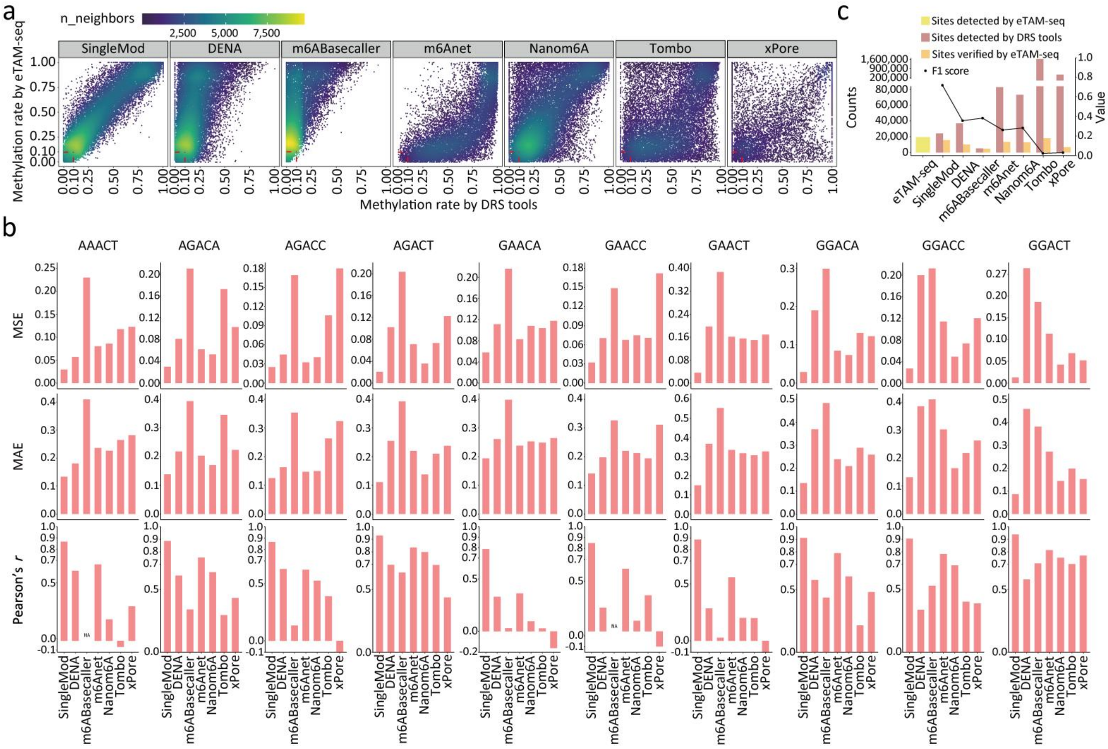

**Supplementary Fig. 4|Comparing the methylation rates in various motifs predicted by DRS tools to eTAM-seq benchmarks on *M. musculus* data. a** Comparing the methylation rates predicted by DRS tools to eTAM-seq benchmarks. **b** The performance of SingleMod and other tools on various motifs. The Pearson's r of m6ABasecaller in certain motifs was designated as "NA", because this tool predicted these motifs in all molecules to be unmethylated. **c** The number of identified m6A sites within the candidate list by DRS tools and eTAM-seq, and their overlap. F1 scores of site-level m6A predictions are also provided.

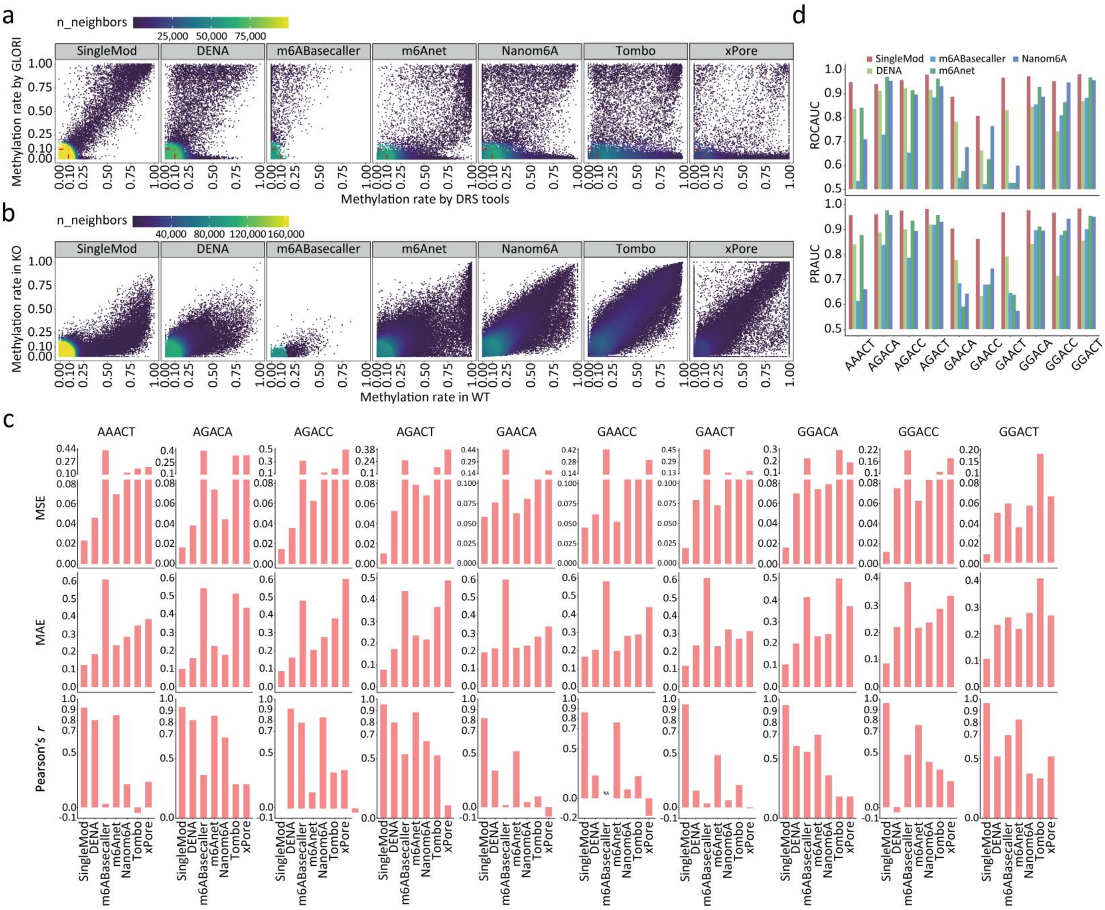

**Supplementary Fig. 5|Comparison of the performance between SingleMod and other tools on *A. thaliana* data. a** Comparing the methylation rates predicted by DRS tools to GLORI benchmarks. **b** Comparing the methylation rates between WT and KO samples predicted by DRS tools. **c** The performance of SingleMod and other tools on various motifs. **d** The AUC of ROC and PR curve for single-molecule m6A predictions in 10 common motifs. Fully-methylated and -unmethylated sites determined by GLORI in both *M. musculus* and *A. thaliana* were used.

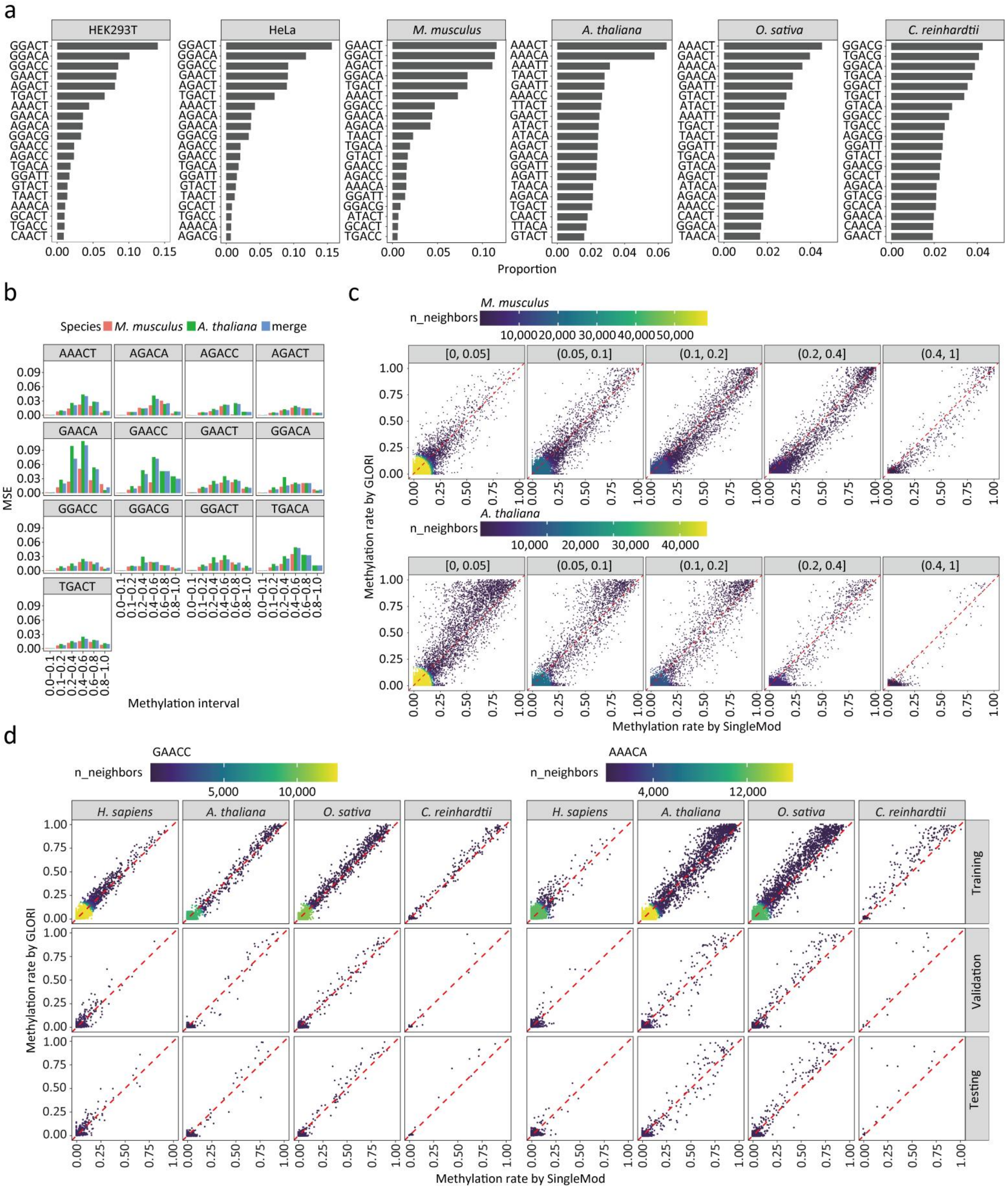


**Supplementary Fig. 6|Further investigation on the performance of SingleMod and optimization of its generalization capacity. a** Top 20 motifs of m6A loacted in multiple species. **b** MSE in varying methylation rate intervals when testing SingleMod in different species. “Merge” indicate merging data from two species. **c** Comparing the methylation rates predicted by SingleMod to GLORI benchmarks for different 9-mers with varying weighted methylation rates in training set. **d** Methylation rate of sites within the GAACC|AAACA motif in different datasets, as detected by GLORI and SingleMod.

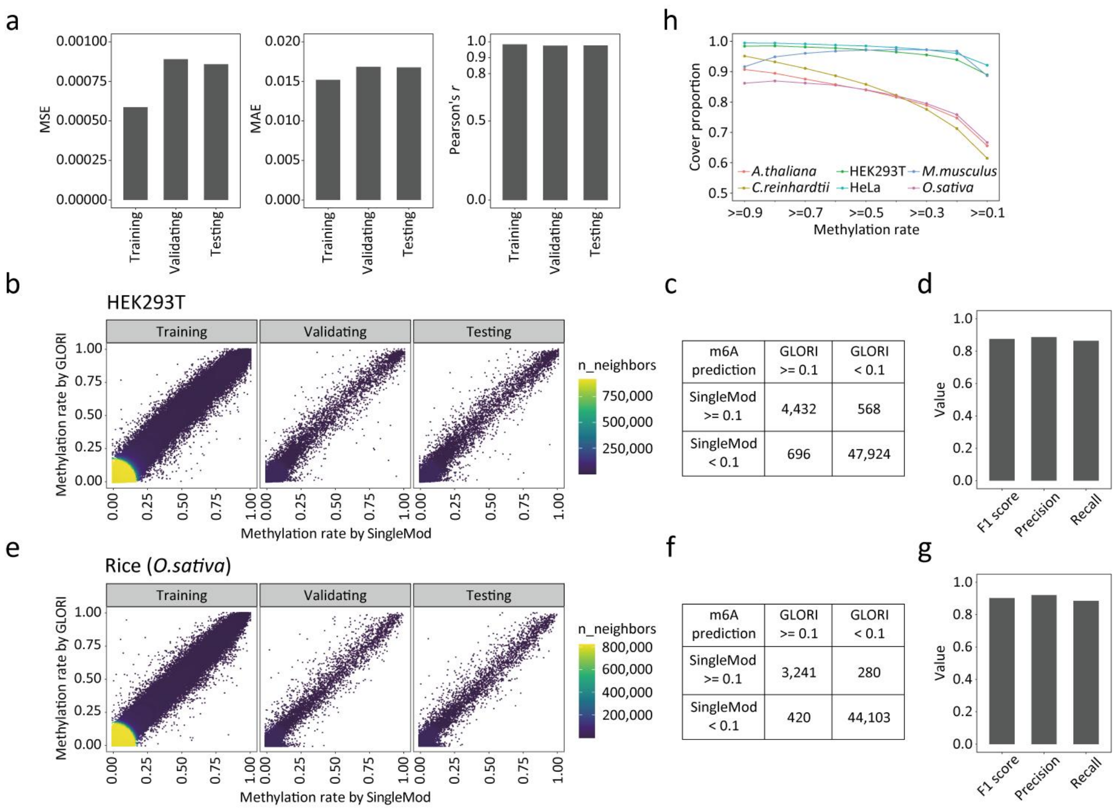


| m6A prediction | GLORI >= 0.1 | GLORI < 0.1 |
|---|---|---|
| SingleMod >= 0.1 | 4,432 | 568 |
| SingleMod < 0.1 | 696 | 47,924 |

| m6A prediction | GLORI >= 0.1 | GLORI < 0.1 |
|---|---|---|
| SingleMod >= 0.1 | 3,241 | 280 |
| SingleMod < 0.1 | 420 | 44,103 |

**Supplementary Fig. 7|Performance of SingleMod trained with DRS data generated by the latest nanopore technology and sequencing kit (RNA004). a** The performance of SingleMod in different datasets. **b** Comparing the methylation rates predicted by SingleMod to GLORI benchmark on different datasets. Data used in **b-d** were from HEK293T. **c** The statistical results of the number of m6A sites with different methylation states predicted by SingleMod and GLORI. Data was from the testing set. **d** F1 scores, precision and recall of site-level m6A predictions by SingleMod. Data was from **c**. **e-g** Results identical to those of **b-d**, but the data came from *O.sativa*.

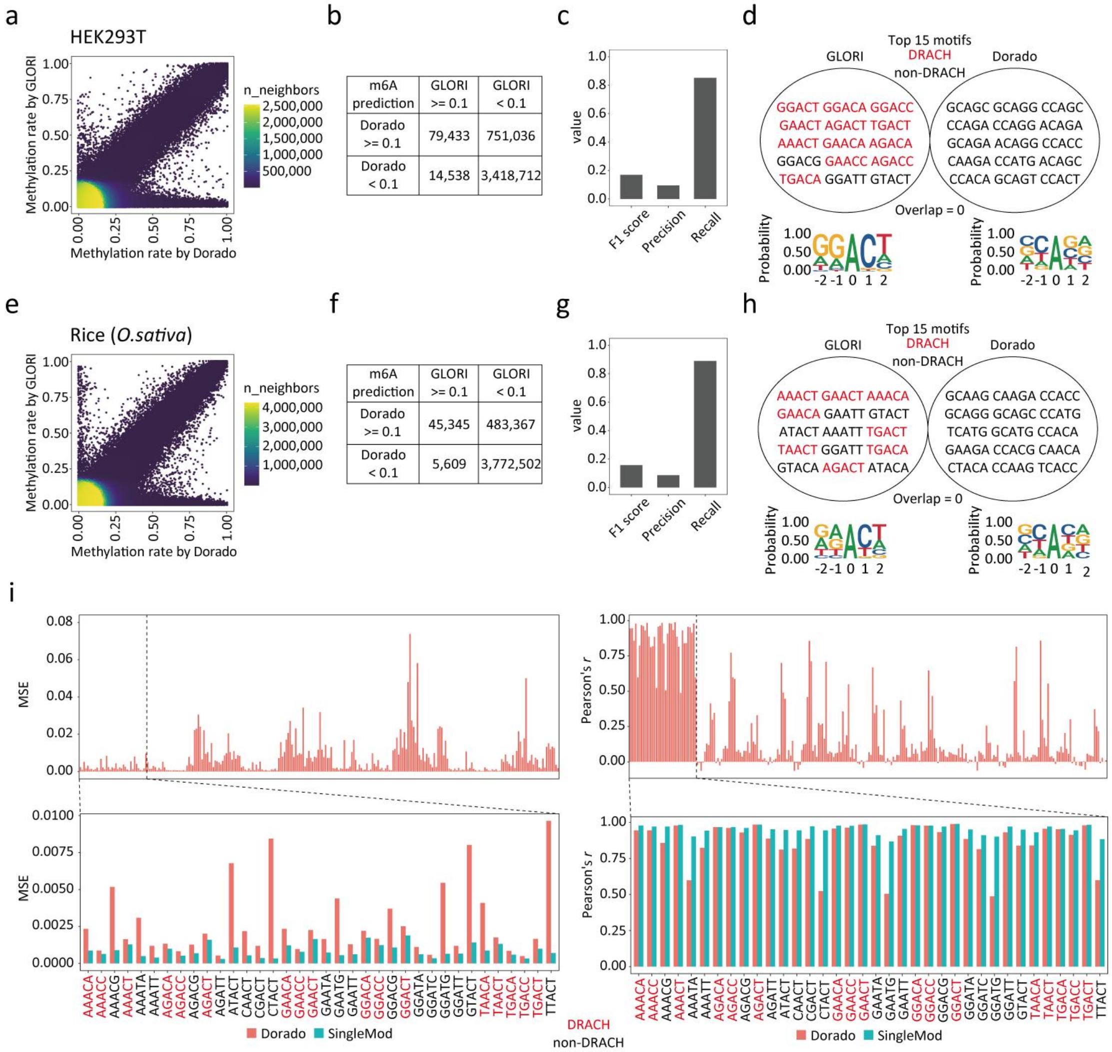


**Supplementary Fig. 8|Performance of Dorado on RNA004 data and comparison with SingleMod (RNA004). a**: Comparing the methylation rates predicted by Dorado to GLORI benchmark on different datasets. Data used in **a-d** were from HEK293T. **b** The statistical results of the number of m6A sites with different methylation states predicted by Dorado and GLORI. **c** F1 scores, precision and recall of site-level m6A predictions by Dorado. **d** Top 15 motifs harboring the most m6A sites identified by GLORI and Dorado. The composition of m6A flanking bases plotted by Seqlogo was shown below. **e-h** Results identical to those of **a-d**, but the data came from *O.sativa*. **i** The performance of Dorado on various motifs, with a focus on the comparison with SingleMod across common motifs.

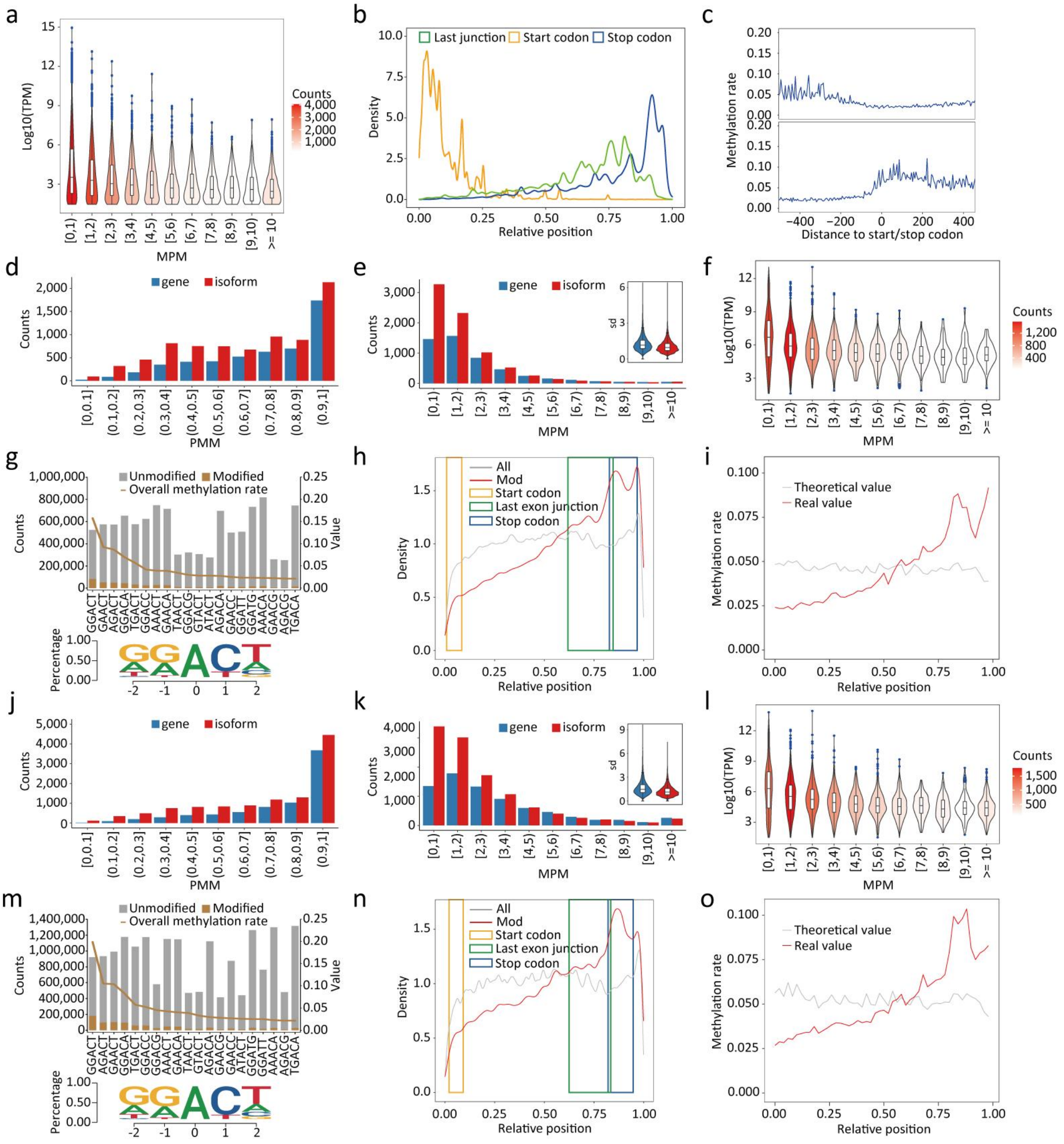

**Supplementary Fig. 9|Single-molecule m6A profiles in human cell lines. a** Distribution of expression levels of isoforms with varying MPM. "TPM" indicates Transcripts Per Million. Data used in **a-c** were from HEK293T. **b** The distribution of relative position of start codon, last junction and stop codon within their molecules. **c** Overall methylation rate around start codon (top) and stop codon (bottom). **d** Counts of genes or isoforms with varying PMM. Data used in **d-i** were from HeLa. **e** Counts of genes or isoforms with varying MPM, and the distribution of standard deviation in m6A number among molecules within each gene or isoform. **f** Distribution of expression levels of genes with varying MPM. **g** Counts (top, bar) of unmodified and modified A bases within top 20 motifs for methylation rate from all molecules and corresponding overall methylation rates (top, line). The composition of m6A flanking bases plotted by Seqlogo are shown below. **h** The distribution of relative position of background (All) and modified (Mod) A bases within their respective molecules. The relative regions of the start codon, last exon junction and stop codon are also marked. **i** Overall methylation rate at different relative positions. **j-o** Results identical to those of **d-i**, but the data came from K562.

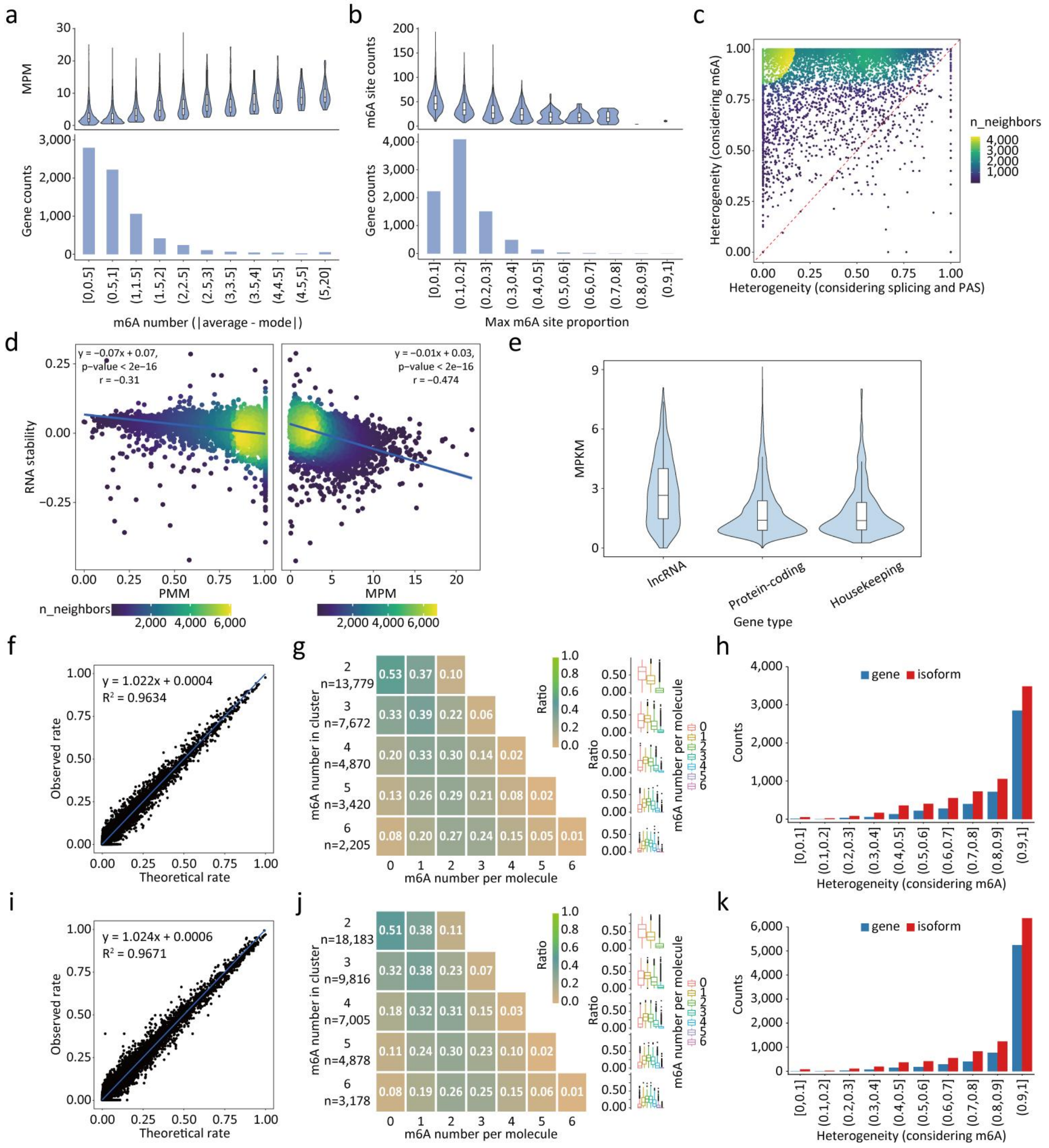


**Supplementary Fig. 10|More information for m6A-mediated RNA heterogeneity and regulation profiled at single-molecule level in human cell lines.** **a** Counts of genes with varying max m6A site proportion (bottom) and distribution of m6A sites counts in corresponding genes (top). Max m6A site proportion indicates the proportion of m6A modifications in the site of the highest methylation rate, relative to the total number of m6A modifications in a gene. Data used **a-e** were from HEK293T. **b** Counts of genes with varying difference between the mode and the mean of m6A number among molecules in a gene. **c** Comparing the molecular heterogeneity introduced by m6A and by difference on splicing form and PAS selection for all genes. **d** Correlation between RNA stability and PMM (left) or MPM (right). **e** Distribution of MPKM from different gene types. **f** Comparison of theoretical rate and observed rate of molecules where two adjacent A bases within all detected motifs are both modified. Data used **f-h** were from HeLa. **g** Heatmap (bottom, left) and boxplot (bottom, right) show the ratio of molecules with varying m6A number within m6A cluster. **h** Gene or isoforms counts with varying molecules heterogeneity introduced by m6A. **i-k** Results identical to those of **f-h**, but the data came from K562.

a

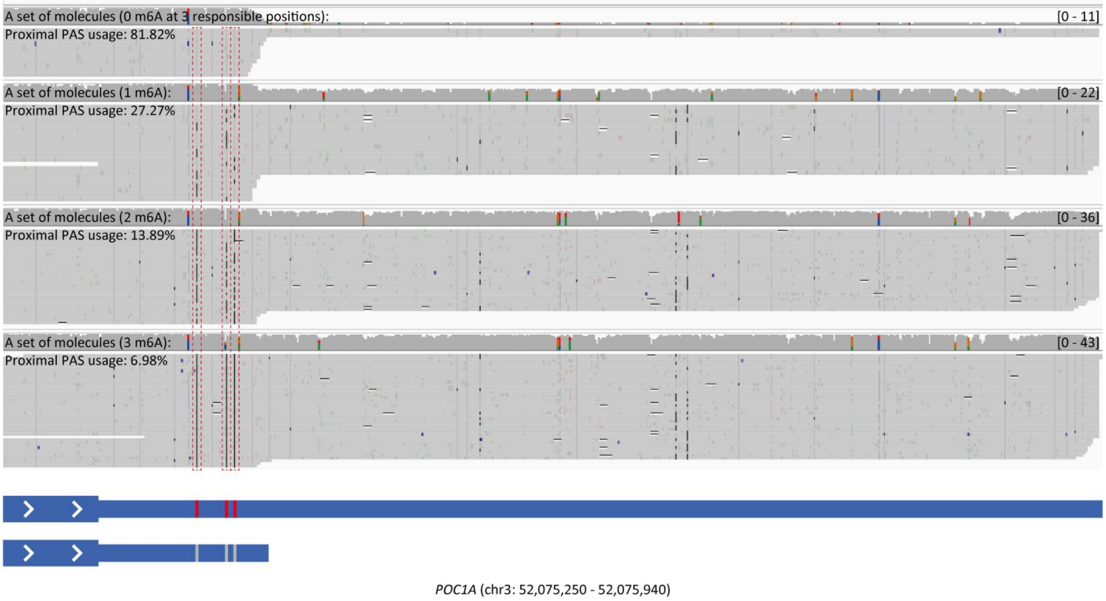


**Supplementary Fig. 11|A significant m6A site-APA correlated events was detected in the gene *POC1A*. In the IGV snapshot which provided a visual representation of this circumstance, reads were grouped based on the modification status of the correlated m6A site (red box) and their corresponding proximal PAS usage are also indicated.**

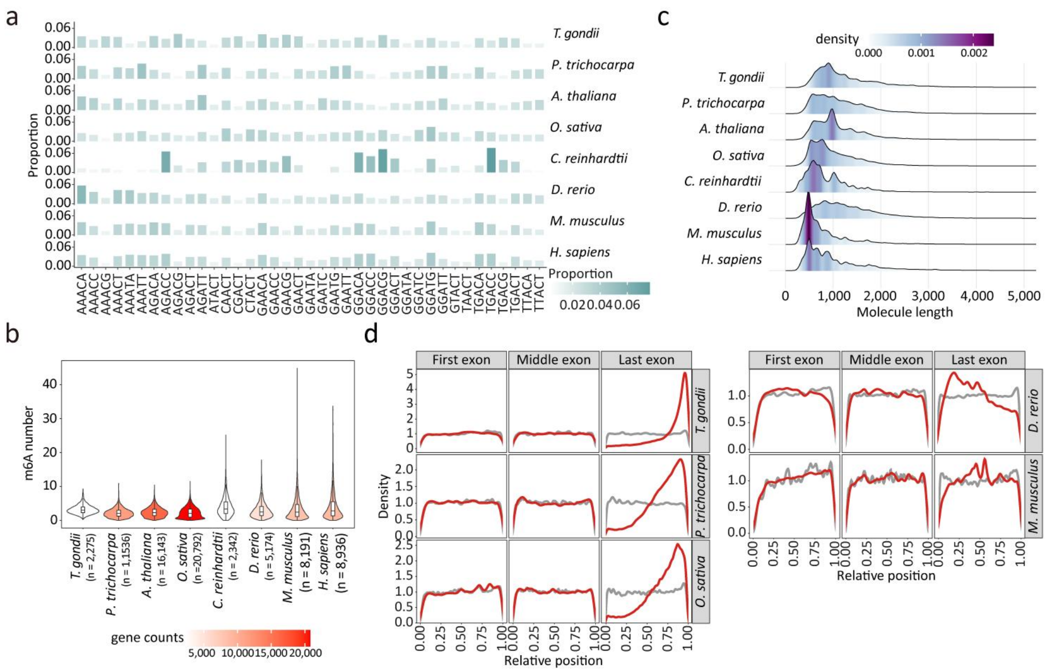

**Supplementary Fig. 12|More information for comparison of single-molecule m6A profiles across multi-species. a** Proportion of candidate A bases within various motifs. **b** Distribution of the average m6A number per molecule for gens. **c** Distribution of molecule length. **d** The distribution of relative position of background (All) and modified (Mod) A bases within their respective molecules in different exons.

a

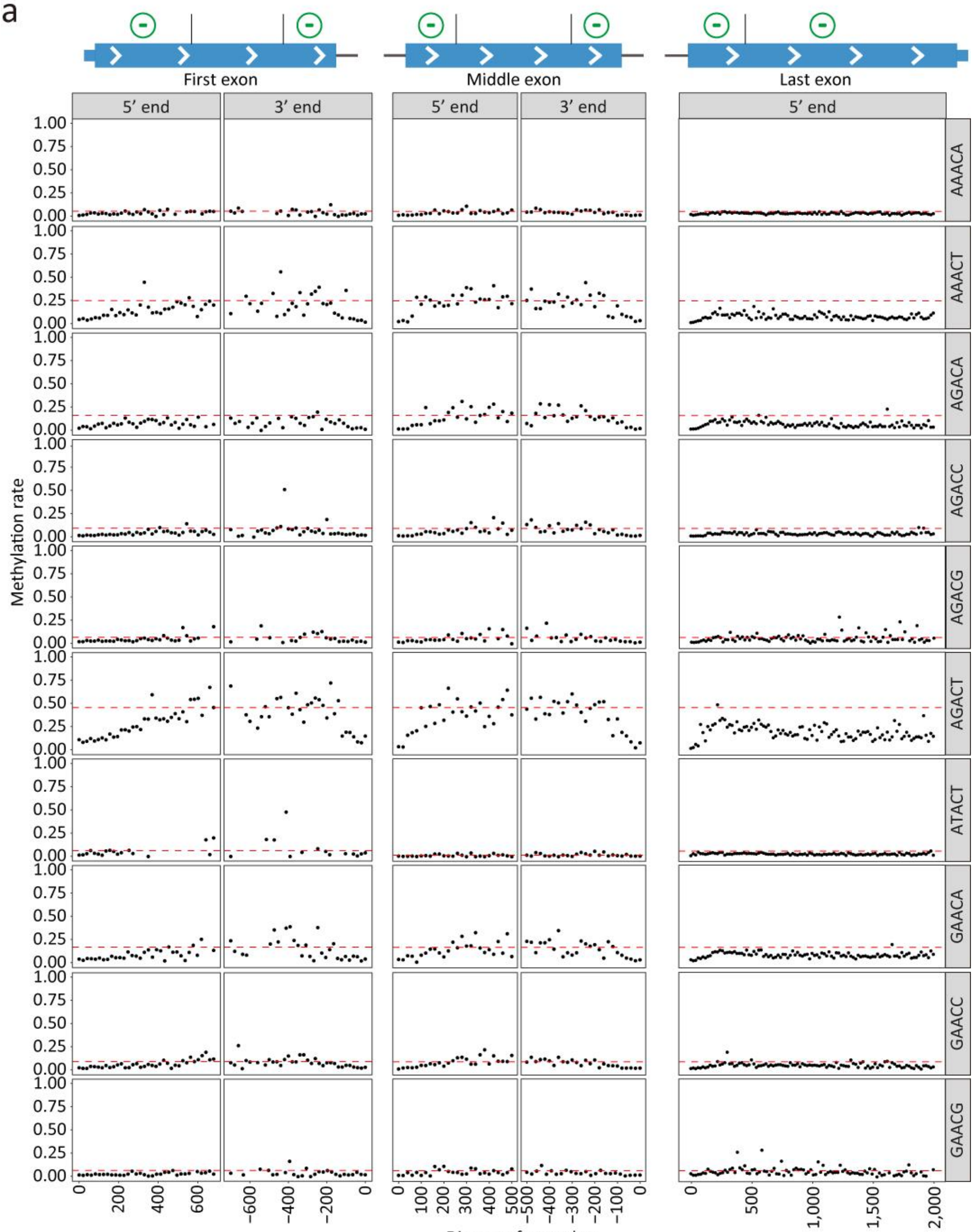

First exon
Middle exon
Last exon
5′ end
3′ end
5′ end
3′ end
5′ end
AAACA
AAACT
AGACA
AGACC
AGACG
AGACT
ATACT
GAACA
GAACC
GAACG
Methylation rate
Distance from edge

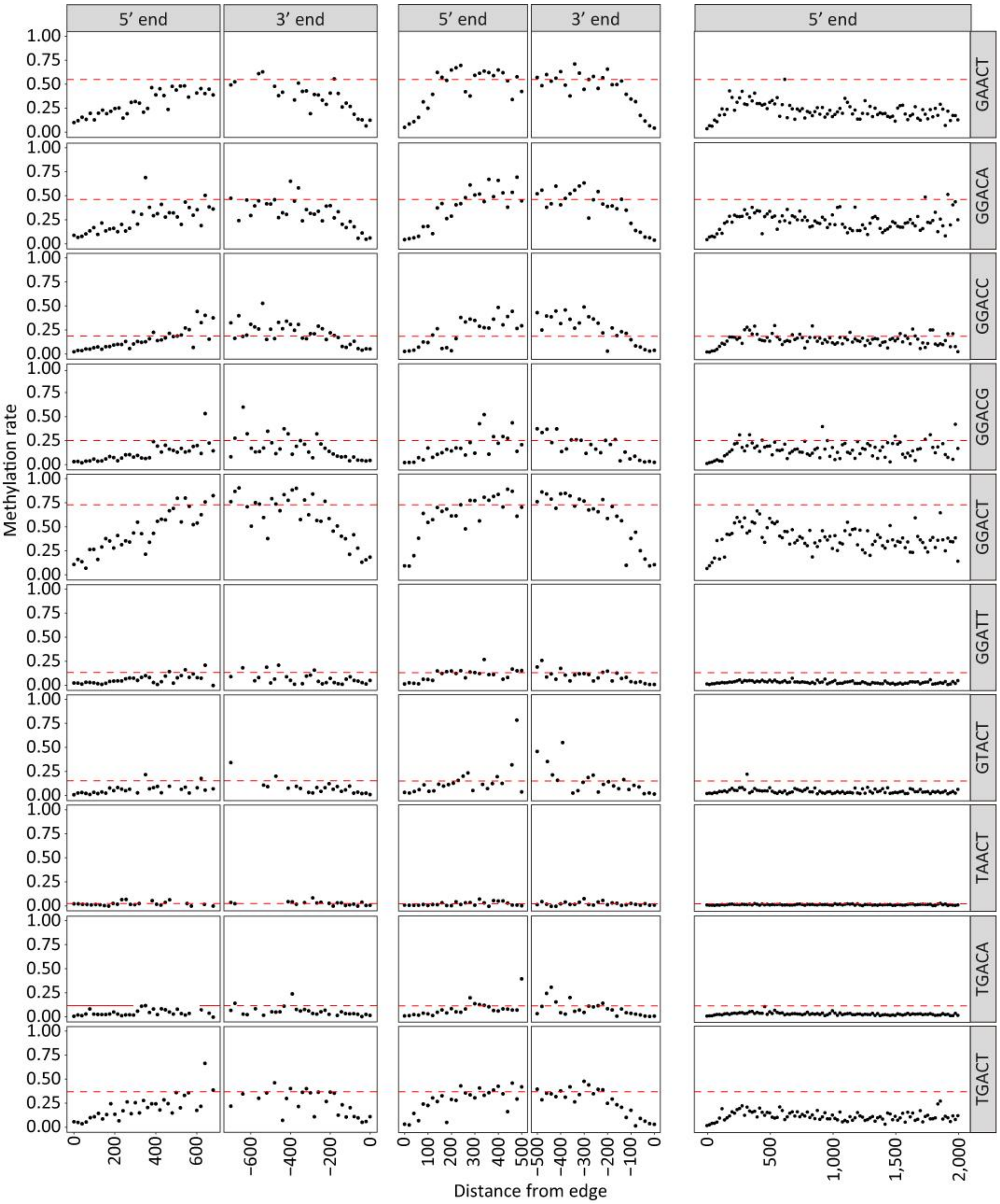


**Supplementary Fig. 13|m6A-pattern shaped by exclusion deposition mode in HEK293T.** Top: schematic representation of exclusion deposition mode. Bottom: m6A pattern shaped by exclusion deposition mode and comparing the overall methylation rate at positions with varying distances from one exon edge to hypothetical methylation rate (red dashed lines).

a

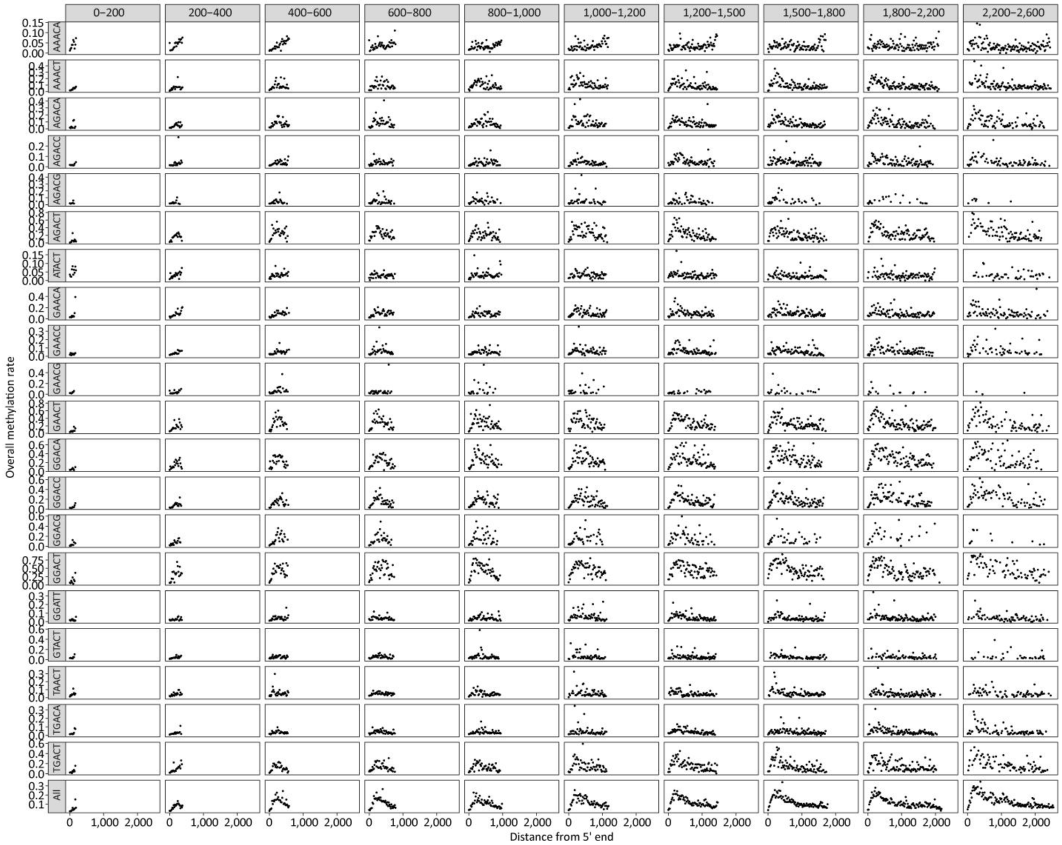


**Supplementary Fig. 14|Overall methylation rate at positions with varying distances from 5' end of last exon with different lengths.** Data are derived from HEK293T.

a

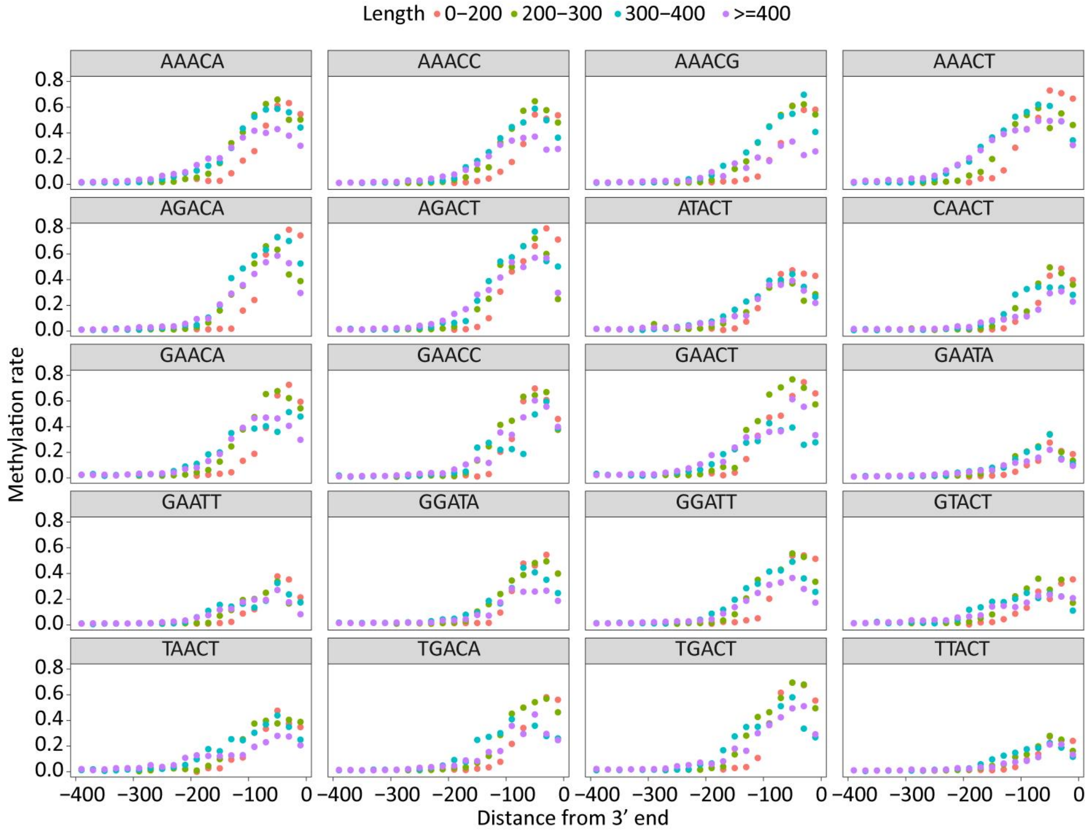


**Supplementary Fig. 15|m6A pattern shaped by inclusion deposition mode and overall methylation rate at positions with varying distances from 3' end of last exon.** Data are derived from *A. thaliana*.

a

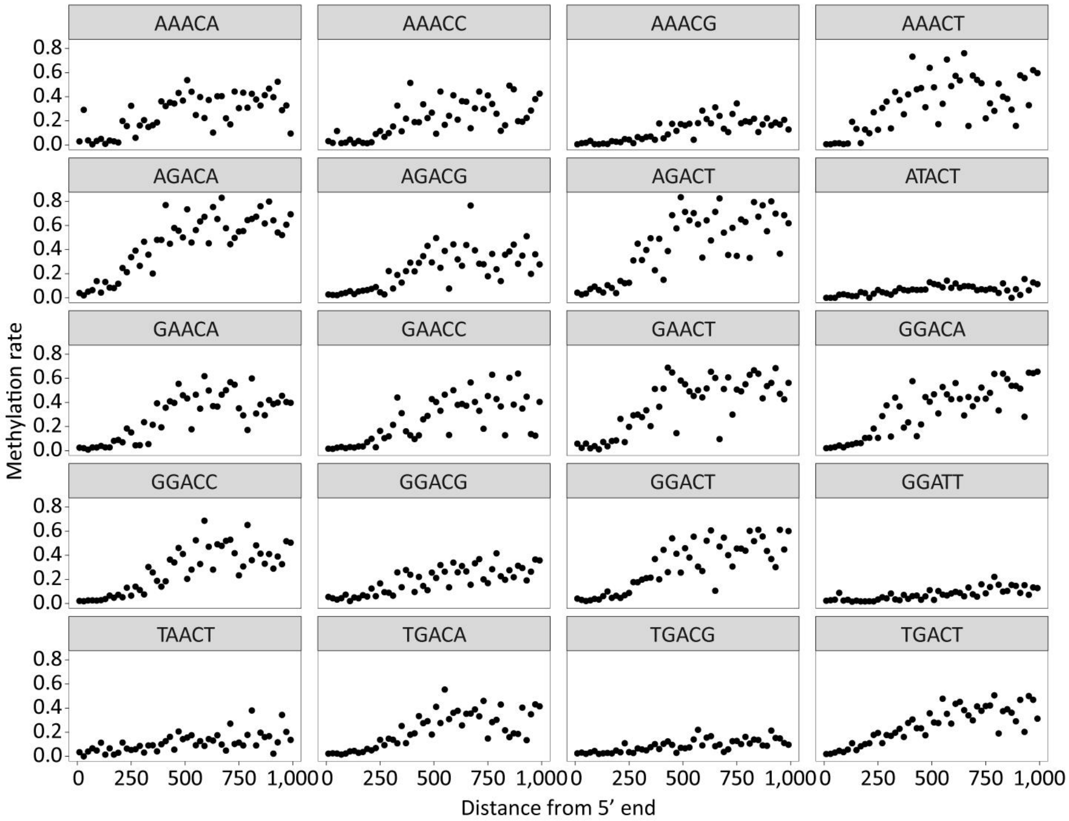


**Supplementary Fig. 16|m6A pattern shaped by combination of inclusion and exclusion deposition mode and overall methylation rate at positions with varying distances from 5' end of last exon.** Data are derived from *C. reinhardtii*.

**Supplementary Tables**

**Supplementary Table 1:** m6A prediction results from mESC DRS data by various tools and their performance as evaluated using GLORI benchmarks.

**Supplementary Table 2:** m6A prediction results from mESC DRS data by various tools and their performance as evaluated using eTAM-seq benchmarks.

**Supplementary Table 3:** m6A prediction results from Arabidopsis DRS data by various tools and their performance as evaluated using GLORI benchmarks.

**Supplementary Table 4:** Evaluation metrics and their values when training final-version and 004-version SingleMod, and A-to-G conversion rate of GLORI experiments.

**Supplementary Table 5:** m6A profiling by SingleMod in various species.

**Supplementary Table 6:** m6A quantification at single-molecule level for different genes and isoforms in human cell lines.

**Supplementary Table 7:** Differential AS and APA events identified in METTL3-KO HEK293T cells and validation results via RT-qPCR in HEK293T cells treated with METTL3 inhibitor.
